# From spikes to intercellular waves: tuning intercellular Ca^2+^ signaling dynamics modulates organ size control

**DOI:** 10.1101/649582

**Authors:** Dharsan K. Soundarrajan, Francisco J. Huizar, Ramezan Paravitorghabeh, Trent Robinett, Jeremiah J. Zartman

## Abstract

Information flow within and between cells depends in part on calcium (Ca^2+^) signaling dynamics. However, the biophysical mechanisms that govern emergent patterns of Ca^2+^ signaling dynamics at the organ level remain elusive. Recent experimental studies in developing *Drosophila* wing imaginal discs demonstrate the emergence of four distinct patterns of Ca^2+^ activity: Ca^2+^ spikes, intercellular Ca^2+^ transients, tissue-level Ca^2+^ waves, and a global “fluttering” state. Here, we used a combination of computational modeling and experimental approaches to identify two different populations of cells within tissues that are connected by gap junctional proteins. We term these two subpopulations “initiator cells” defined by elevated levels of Phospholipase C (PLC) activity and “standby cells,” which exhibit baseline activity. We found that the strength of hormonal stimulation and extent of gap junctional communication jointly determine the predominate class of Ca^2+^ signaling activity. Further, single-cell Ca^2+^ spikes are stimulated by insulin, while intercellular Ca^2+^ waves depend on Gαq activity. Our computational model successfully recapitulates how the dynamics of Ca^2+^ transients varies during organ growth. Phenotypic analysis of perturbations to Gαq and insulin signaling support an integrated model of cytoplasmic Ca^2+^ as a dynamic reporter of overall tissue growth. Further, we show that perturbations to Ca^2+^signaling tune the final size of organs. This work provides a platform to further study how organ size regulation emerges from the crosstalk between biochemical growth signals and heterogeneous cell signaling states.

**Author Summary:** Calcium (Ca^2+^) is a universal second messenger that regulates a myriad of cellular processes such as cell division, cell proliferation and apoptosis. Multiple patterns of Ca^2+^ signaling including single cell spikes, multicellular Ca^2+^ transients, large-scale Ca^2+^ waves, and global “fluttering” have been observed in epithelial systems during organ development. Key molecular players and biophysical mechanisms involved in formation of these patterns during organ development are not well understood. In this work, we developed a generalized multicellular model of Ca^2+^ that captures all the key categories of Ca^2+^ activity as a function of key hormonal signals. Integration of model predictions and experiments reveals two subclasses of cell populations and demonstrates that Ca^2+^ signaling activity at the organ scale is defined by a general decrease in gap junction communication as organ growth. Our experiments also reveal that a “goldilocks zone” of optimal Ca^2+^ activity is required to achieve optimal growth at the organ level.

## Introduction

Epithelial cells do not observe social distancing. This is evident during epithelial morphogenesis when cells communicate and coordinate their activities to generate functioning organs (Leptin, 1995; Schöck and Perrimon, 2002). One modality of intercellular communication occurs through gap junctions (GJ), intercellular channels that permit direct cell-cell transfer of ions and other small molecules (Levin, 2007). Calcium ions (Ca^2+^) act as second messengers that regulate a myriad of cellular processes such as proliferation, differentiation, transcription, metabolism, cellular motility, fertilization, and neuronal communication (Berridge, 2005; Carafoli, 2002; Clapham, 2007; Cuthbertson and Cobbold, 1985; Giorgi et al., 2018; Humeau et al., 2017; La Rovere et al., 2016; Martin and Bernard, 2017; Orrenius et al., 2003; Wei et al., 2009). Ca^2+^ signaling also regulates developmental processes at the multicellular level. For instance, Ca^2+^ signaling has been shown to regulate scale development in butterfly wings (Ohno and Otaki, 2015). It also mediates autophagic and apoptotic processes required for hearing acquisition in the developing cochlea (Ceriani et al., 2016; Mammano and Bortolozzi, 2018; Takeuchi et al., 2020). However, a systems-level description of Ca^2+^ signaling during organ development is lacking.

A major challenge in reverse engineering Ca^2+^ signaling during organ development is the lack of an in vivo model system to identify how cells interpret and integrate information across the broad range of input molecules that dynamically vary concentrations of cytosolic Ca^2+^ ions. In particular, it remains unclear how single-cell Ca^2+^ dynamics are coordinated to regulate tissue-level Ca^2+^ patterns. To overcome these challenges, we developed a computational model based on a realistic geometry of epithelial cells to model Ca^2+^ signaling in the *Drosophila* wing imaginal disc. The *Drosophila* wing imaginal disc is an experimentally amenable system for investigating systems-level regulation of cell signaling (Figure 1A) (Balaji et al., 2017; Brodskiy et al., 2019; Restrepo and Basler, 2016). *Drosophila* wing imaginal discs are a premier system to gain insights into several organ-intrinsic and organ-extrinsic mechanisms that control organ growth (Buchmann et al., 2014; Gou et al., 2020; Hariharan, 2015; Restrepo et al., 2014; Vollmer and Iber, 2016; Vollmer et al., 2017).

**Figure 1:**
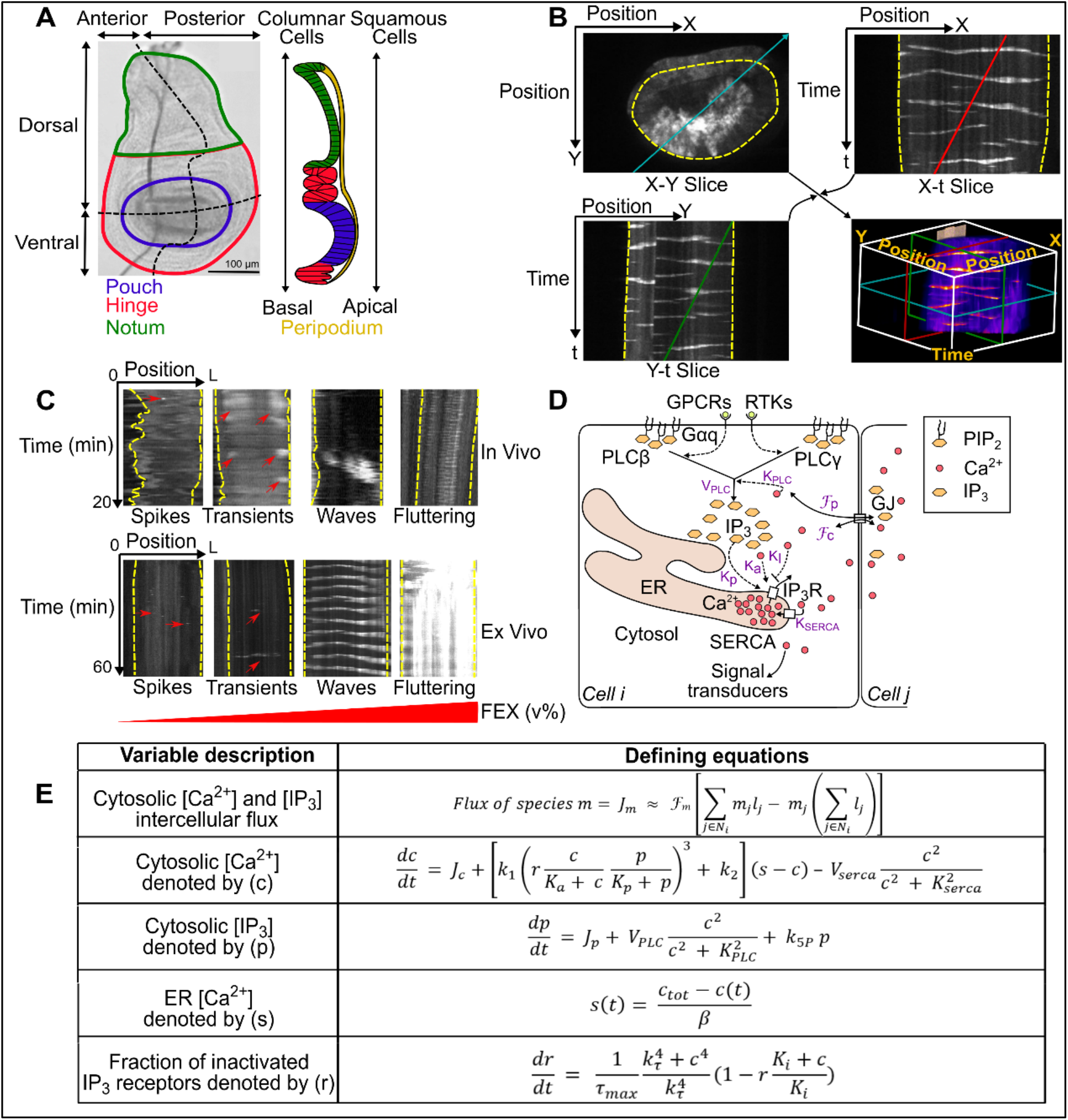
Multicellular Ca^2+^ signaling in a developing organ. **(A)** (*Left panel*) Image of third instar *Drosophila* wing imaginal disc. The larval wing disc includes four regions: pouch (blue), hinge (red), notum (green) and peripodium (yellow). The pouch cells are the region of interest for this study. (*Right panel*) A schematic of the side view of the wing disc showing the peripodial membrane composed of squamous epithelial cells. **(B)** Kymographs illustrate two-dimensional slices of three-dimensional (X, Y and t planes respectively) spatiotemporal signaling. Ca^2+^ signaling activity is related to the fluorescence intensity of the GCaMP6f sensor. A line in the X-Y plane (blue), the X-t plane (red), and Y-t plane (green) are overlaid to illustrate the positions of 2D projections in 3D data set. **(C)** Four classes of Ca^2+^ signaling patterns are observed both in vivo and ex vivo: single-cell spikes, intercellular transient, intercellular waves and fluttering (Brodskiy et al., 2019). In ex vivo cultures, the occurrence of these patterns depend on the concentration of fly extract added to the culture media. Red arrows highlight subtle cellular Ca^2+^ activity. **(D)** Major components of Ca^2+^ toolkit: G protein–coupled receptors (GPCR), receptor tyrosine kinase (RTK), gap junctions (GJ), Inositol trisphosphate (IP_3_), diacylglycerol (DAG), Phospholipase C (PLC), Phosphatidylinositol 4,5-bisphosphate (PIP_2_), sarco/endoplasmic reticulum Ca^2+^-ATPase (SERCA) and IP_3_ receptors (IP_3_R). Parameters for our computational model are denoted in purple. K_PLC_ values are presumably a function of the specific PLC isoforms active in the cell. **(E)** A summary of baseline Ca^2+^ signaling model. More details on each dynamical equation and meaning of the parameters in the model are described in Supporting Information and text.

Previous experimental investigations revealed four classes of Ca^2+^ signaling activity in the developing wing disc: single-cell Ca^2+^ spikes, multicellular transients, intercellular waves, and global fluttering (Figure 1B, C). Recently we showed that these patterns depend on the strength of agonist stimulation (Brodskiy et al., 2019). We and others have previously reported that Ca^2+^ patterns observed in the wing disc are dependent on phospholipase C (PLC) and the inositol trisphosphate receptor (IP_3_R) pathway mediated by gap junctional communication (Balaji et al., 2017; Brodskiy et al., 2019; Restrepo and Basler, 2016). In non-excitable cells, stimulation of receptors in the cell surface results in activation of PLCs to generate IP_3_, which binds to and activates IP_3_R (Figure 1D) (Clapham, 2007). Upon binding, IP_3_Rs channel Ca^2+^ from the endoplasmic reticulum (ER) to the cytosolic space (Berridge, 1987; Berridge et al., 2000; Clapham, 2007). However, the specific receptors involved in stimulation of PLCs in the *Drosophila* wing imaginal disc remain to be more fully defined. The key physical/chemical parameters and their interactions that define multicellular Ca^2+^ dynamics in response to agonist stimulation is not fully characterized. How these different spatiotemporal modes of signaling encode information from upstream signals to impact downstream cellular processes during organ development is not characterized. We have previously shown that inhibition of Ca^2+^ regulators IP_3_R, PLC21C, Gαq and gap junctional protein innexin 2 (Inx2) in the wing disc results in reduction in size of adult wing blade (Brodskiy et al., 2019). Whether changes to multiscale Ca^2+^ signaling patterns in wing disc alters overall adult blade wing size remains unknown.

Overall, the computational models of calcium signaling in developing epithelial systems have to date received sparse attention. Here, we report the necessary conditions to generate the full spectrum of experimentally observed spatiotemporal patterns by employing a computational modeling approach. To do so, we built a geometrically accurate 2D-model of a wing disc based on experimental data. We discovered that in silico replication of wing disc Ca^2+^ patterns requires two distinct classes of cells, which we term as “initiator cells” and “standby cells.” We show how the standby cells organize themselves with respect to a V_PLC_ Hopf bifurcation threshold, and how the range of standby cell V_PLC_ values determine the final patterns of Ca^2+^ signaling. Next, computational simulations and experiments demonstrate that gap junctional communication alters Ca^2+^ signaling response resulting in more Ca^2+^ spikes in the absence of external stimuli. Finally, we provide computational and experimental evidence for the role of Ca^2+^ signaling in imaginal disc morphogenesis. Our findings suggest a “goldilocks zone” of Ca^2+^ where lower levels of Ca^2+^ is correlated with reduced organ growth and higher levels of Ca^2+^ is also correlated with reduced growth dependent upon the stimulus triggering the Ca^2+^ signal. Overall, we identify crucial crosstalk between biochemical growth signals, such as insulin and Gαq, and heterogeneous cell signaling states during the growth of an organ.

## Results

### Relative rate of IP_3_ production governs transitions between classes of spatiotemporal Ca^2+^ patterns at the tissue level

Multiple spatiotemporal classes of Ca^2+^ activity are observed in vivo and ex vivo in the wing disc. However, an understanding of how this activity is regulated requires developing a systems-level description. To summarize, these include: (i) single-cell Ca^2+^ spikes, (ii) intercellular Ca^2+^ transients (ICTs), (iii) intercellular Ca^2+^ waves (ICWs), and a (iv) global fluttering phenomenon (Figure 1C, Table S1, SI videos 1-8) (Balaji et al., 2017; Brodskiy et al., 2019; Restrepo and Basler, 2016). The frequencies of these observed classes are dependent on the age of the larvae in both in vivo and ex vivo experiments. Younger larva with smaller discs (4-5 days after egg laying) exhibit higher occurrences of ICWs and fluttering states while older, larger larval discs (6-8 days after egg laying) predominantly show ICTs and spikes (Brodskiy et al., 2019). For ex vivo cultures, the transition from limited to tissue-wide calcium activity depends on the amount of fly extract (FEX) added to the culture. Low concentrations of FEX stimulated Ca^2+^ spikes. Progressively increasing levels of FEX resulted in ICTs, ICWs, and eventually fluttering. Further, FEX-stimulated Ca^2+^ dynamics is based on IP_3_R-based release of Ca^2+^ from the ER to the cytosol as shown in Figure 1D (Brodskiy et al., 2019).

These experimental findings motivated us to ask what specific cellular properties of the wing disc result in the emergence of these distinct patterns. To systematically investigate the underlying principles governing the emergence of these patterns, we formulated a two-dimensional image-based geometrically realistic computational model of Ca^2+^ signaling in the wing disc pouch where the columnar epithelial cells are connected by gap junctions (Figure S1), (Jursnich et al., 1990; Ryerse, 1982; Weir and Lo, 1982, 1985). Image-based modelling enables the wholistic characterization of molecular mechanisms and tissue dynamics during organogenesis (Gómez et al., 2017). Given the near universal conservation of the Ca^2+^ signaling pathway across model systems (Berridge, 1987; Berridge et al., 2000), the baseline single-cell Ca^2+^ model in our study was adapted from Politi and colleagues (Politi et al., 2006). The model equations and the descriptions of the parameters are shown in Figure 1 D, E and Table S1. We modified the rate of IP_3_R inactivation term, *r*, to replicate our experimental Ca^2+^ dynamics. The modified equation is described below:

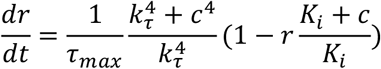

A similar modification to the rate of inactivation term, *r*, has been proposed previously (Sneyd et al., 2017).

To reproduce the four distinct patterns in silico, we varied the V_PLC_ parameter progressively in individual cells across a range of parameters (Table S1). Given that our patterns were dependent on the concentration of FEX, we varied V_PLC_ as a lumped-parameter representing the level of agonist stimulation. The computational model successfully reproduced the four different spatiotemporal classes of Ca^2+^ signaling dynamics observed in vivo and ex vivo (Figure 2A-D, SI videos 9-12). Interestingly, we discovered that the formation of these patterns is dependent on the number of cells having a V_PLC_ value below, above, or equal to the Hopf bifurcation threshold for single-cells (V_PLC_ = 0.774) (Figure 2E). Simulated cells that have a V_PLC_ value above the Hopf threshold are termed “initiator cells” and are posed to exhibit high levels of IP_3_ production. Neighboring simulated cells with V_PLC_ values below the Hopf threshold are termed “standby cells” that receive a signal from initiator cells to propagate a signal. For instance, if a majority of standby cells have V_PLC_ values significantly below the critical Hopf bifurcation threshold, single-cell Ca^2+^ spikes occur only where initiator cells are present (Figure 2A, E). When we increased standby cell V_PLC_ values close to the lower end of the Hopf bifurcation point (Figure 2B, E), we noticed the formation of ICTs. Finally, we observed the formation of ICWs and fluttering phenotypes (Figure 2C, D, E) for cases when the majority of cells in the system were assigned a V_PLC_ above the bifurcation threshold, thereby placing most cells in an initiator state. In the absence of initiator cells, Ca^2+^ activity is not observed. This suggests that initiator cells are necessary for the formation of Ca^2+^ transients in developing tissues.

**Figure 2:**
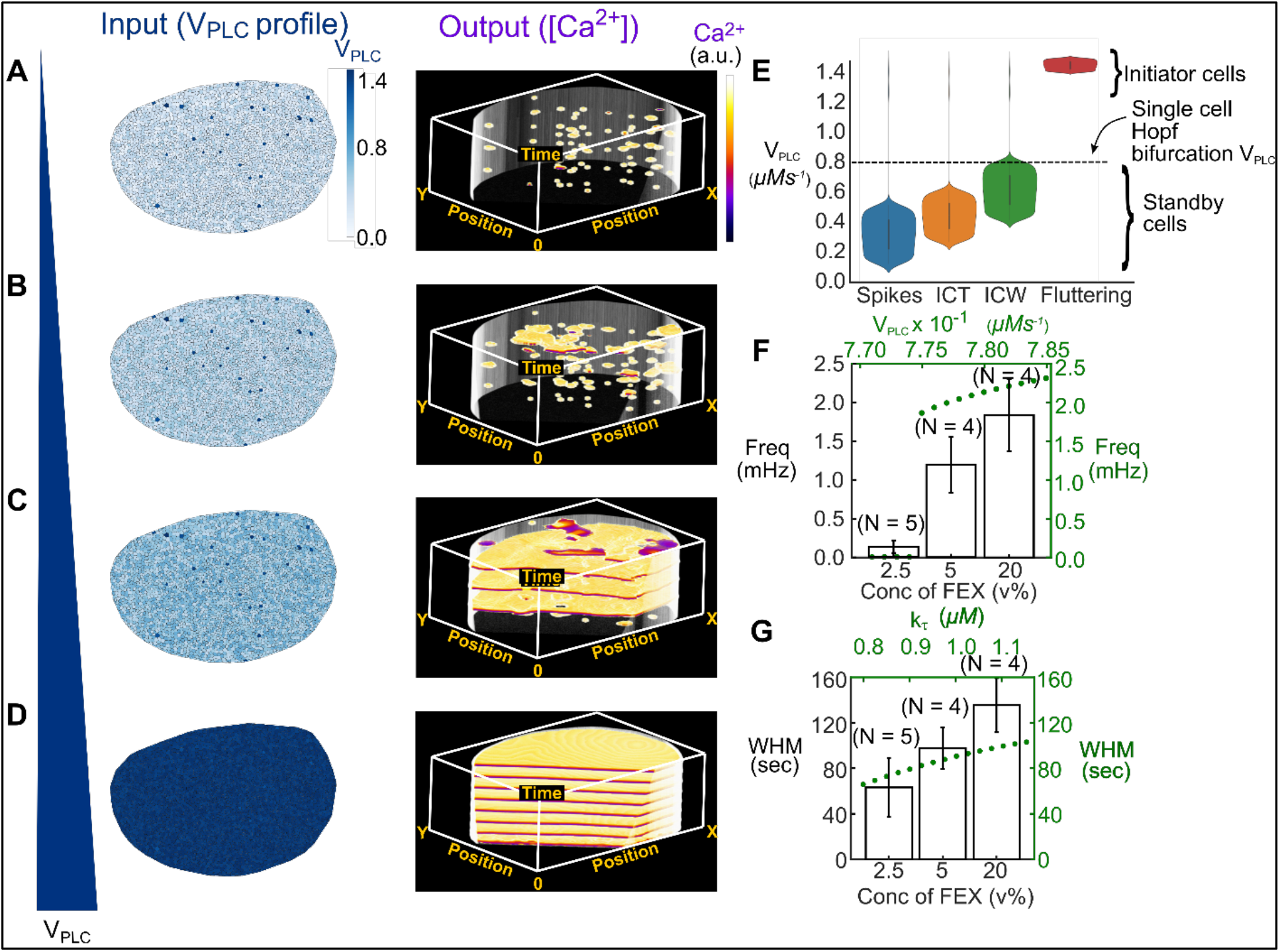
A progression of hormonal stimulation governs the spatial extent of intercellular Ca^2+^ communication. **(A-D)** Computer simulations recapitulating the key classes of multicellular Ca^2+^ activity observed in vivo and ex vivo. **(A)** When the majority of cells have V_PLC_ values below the Hopf bifurcation threshold (*left*), single-cell Ca^2+^ spikes are seen (*right*). **(B)** Intercellular Ca^2+^ transients are observed (*right*)as the distribution of V_PLC_ in cells is increased (*left*). **(C)** A further increase in V_PLC_ (*left*) results in the emergence of periodic intercellular Ca^2+^ waves (*right*). **(D)** “Fluttering” occurs (*right*) when V_PLC_ levels in all of the cells in the disc is above Hopf bifurcation (*left*). **(E)** Quantification of V_PLC_ distribution in the initiator and the standby cells for each of the classes of Ca^2+^ signaling activity. The dashed line indicates the region in V_PLC_ parameter space permitting Ca^2+^ oscillations. **(F-G)** The single-cell version of our model predicts that the frequency and width at half maximum (WHM) of Ca^2+^ oscillations is altered by varying V_PLC_ and *k_τ_*.This prediction matches the WHM of Ca^2+^ activity observed in ex vivo discs cultured with variable concentrations of fly extract. Error bars are reported as standard error of the means (SEM).

The “single-cell” version of the model predicts that differences in Ca^2+^ signal amplitude and frequency are tunable by varying the V_PLC_ and *k_τ_* parameters (Figure S2 A,B). To further examine the influence of these and other parameters on Ca^2+^ signaling dynamics, two-dimensional computational model simulations of the tissue were performed by varying V_PLC_, *k_τ_* and *τ_max_* (Figure S3B-D) as a percentage of the baseline signal while holding other parameters constant. The baseline parameter set was selected such that the simulation resulted in intercellular Ca^2+^ waves (Figure S3, red box). Reducing *k_τ_* leads to a narrower width at half maximum (WHM) of Ca^2+^ transients (Figure S3B). A similar result is observed when a reduction of *τ_max_* results in a decrease in WHM, whereas an increase in *τ_max_* results in an increase in WHM (Figure S3C). A decrease in the *τ_max_* parameter increases the frequency of signals in the simulated tissue, while an increase reduces the frequency of Ca^2+^ transients. This suggests that the system is more sensitive to *τ_max_*, and variations in *τ_max_* have much greater impact in how quickly the system responds to stimulus.

These results indicate that tissue-level Ca^2+^ patterns depend on the spatial distribution of cell states defined by their maximal IP_3_ production rates, in relation to the effective tissue-level Hopf bifurcation threshold. Further, key parameters in the model can be modified in a manner to allow tunable Ca^2+^ signaling patterns to influence signal strength, duration, and propagation.

### GJ communication limits Ca^2+^ spikes in the absence of hormonal stimulation

To elucidate how gap junctional proteins alter Ca^2+^ signals, we simulated a scenario where the initiator cells chosen at random had their V_PLC_ values set to the Hopf bifurcation threshold value of 0.774 and standby cells had a V_PLC_ values that were randomly uniformly distributed between 0.1 and 0.5 (Figure 3A). Under these conditions, no Ca^2+^ activity was observed in the presence of normal functioning GJ communication (Figure 3A’). Next, we compared the effect of blocking gap junctional communication in silico. To do so, we set the permeability terms for Ca^2+^ (F_c_) and for IP_3_ (F_p_) to zero. Strikingly, we observed Ca^2+^ spikes in simulated wing disc cells in the absence of GJ communication (Figure 3A”). We explored this phenomenon computationally by considering a single stimulated cell connected to neighboring cells by GJ communication by performing bifurcation analysis on our modified model for a single-cell. We observed the emergence of a Hopf bifurcation as expected (Figure S4A). Next, the effect of the initial Hopf bifurcation point (HB_1_) on gap junctional permeability of IP_3_, F_p_, was analyzed. Setting F_c_ to zero and progressively increasing F_p_ increased the critical maximum rate of IP_3_ activation threshold 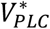 where HB_1_ occurred (Figure S4B). Similar trends were observed when F_c_ was increased to 0.05.

**Figure 3:**
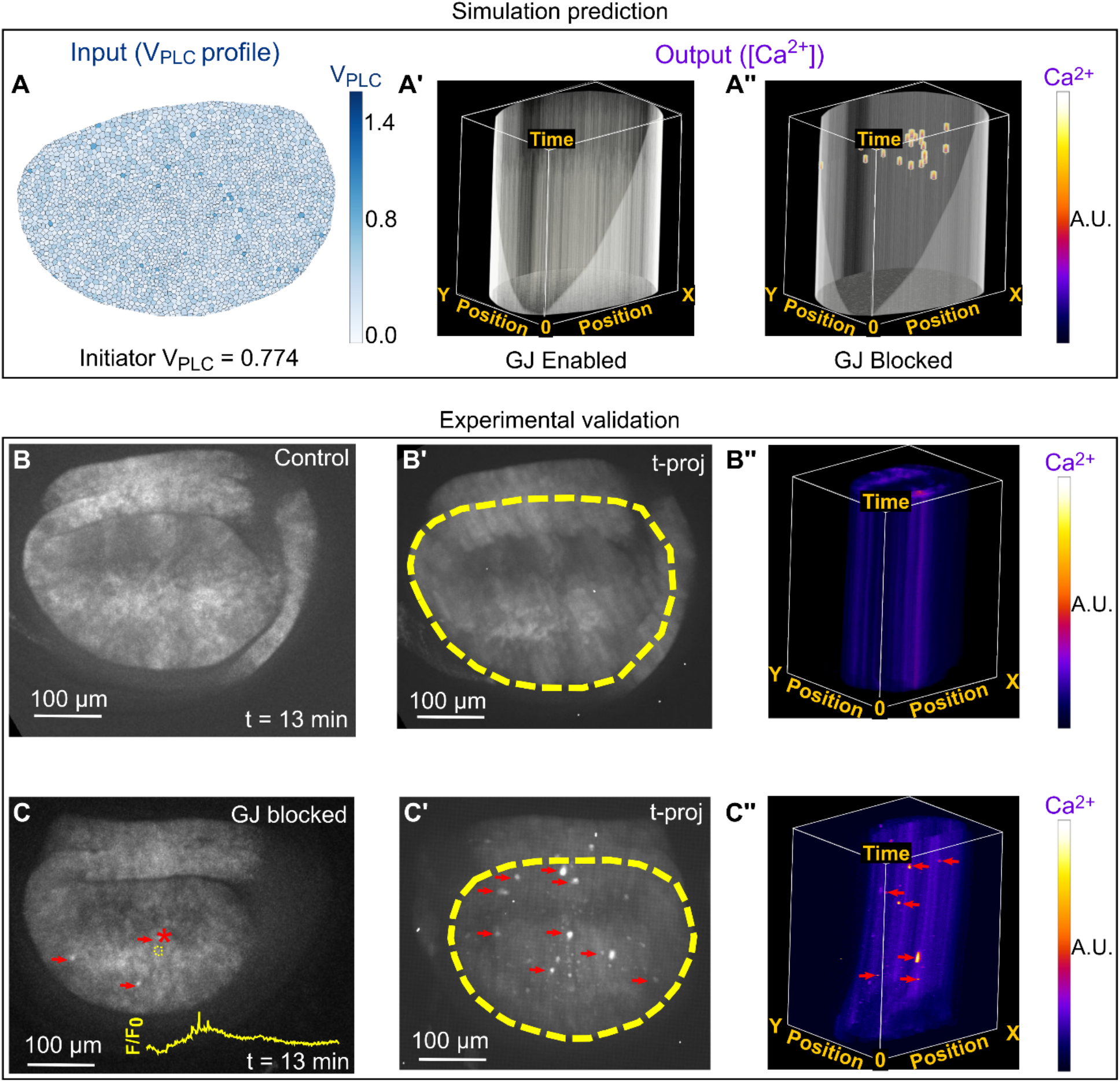
Gap junction (GJ) communication decreases the proportion of cells exhibiting Ca^2+^ spikes. **(A)** Simulation of Ca^2+^ signaling in wing disc where the V_PLC_ values of initiator cells were set to the Hopf bifurcation threshold 0.774, and standby cell V_PLC_ values were randomly distributed between 0.1 and 0.5. **(A’)** Allowing GJ communication by letting permeability of IP_3_ (F_p_) and Ca^2+^ (F_c_) be 0.005 and 0.0005 respectively, resulted in no Ca^2+^ activity (GJ Enabled). F_p_ and F_c_ were set to zero to simulate inhibition of GJ communication (GJ Blocked). Inhibition of GJ communication resulted in Ca^2+^ spike activity. **(B)** Ex vivo time lapses of nub>GCaMP6f x UAS-RyR^RNAi^ (control) wing discs in Grace’s low ecdysone media were generated by imaging for 1 h at 10 sec intervals. Under this condition, we observed no Ca^2+^ activity in the wing disc cells. **(B’)** Time-projection of the time lapse stack. The wing disc boundary is indicated with the yellow dashed line. **(B”)** A kymograph generated further demonstrates no instances of Ca^2+^ activity. **(C)** GJ communication was blocked by culturing wing discs in Grace’s low ecdysone media with 100 mM of Carbenoxolone (CBX). Instances of spike activity are denoted by red arrows. The intensity of a region of interest (yellow dashed circle) is overlaid to demonstrate a spike in local Ca^2+^ activity. Ca^2+^ spike is observed when the intensity normalized to basal intensity is plotted (yellow line, F/F_0_). **(C’)** Time-projection of the time lapse movies. We observed a significant number of Ca^2+^ spikes in the 1 h time interval when GJs were inhibited. Yellow dashed lines indicated disc boundary. **(C’’)** A kymograph generated demonstrates instances of Ca^2+^ spike activity.

A closer look into the importance of GJ permeability on the formation of Ca^2+^ signals was done by varying F_c_ and F_p_ in computational simulations. Starting from a baseline ICW, a decrease in GJ permeability resulted in a transition from ICWs to ICTs, and eventually to single-cell Ca^2+^ spikes, while an increase in GJ permeability did not perturb the baseline signal (Figure S3A). These findings show that gap junctional permeability alters the cytosolic residence time for critical messengers such as IP_3_ and Ca^2+^ whose cytosolic concentrations affects Ca^2+^ release from ER in both initiator cells and standby cells. This is consistent with bifurcation analysis demonstrating that gap junctional permeability influences stimulation threshold required for Ca^2+^ oscillations in cells (Figure S4B).

We tested these computational modeling predictions experimentally. To do so, we pharmacologically inhibited the gap junctional protein Inx2 using Carbenoxolone (CBX) and observed the emergence of Ca^2+^ spikes in the absence of FEX in the culture media (Figure 3B-B”, C-C”, SI movies 13,14). This further demonstrates that gap junction communication regulates Ca^2+^ dynamics in the wing disc pouch.

### Gap junction permeability shapes Ca^2+^ signaling during development

Of note, integrated intensity of Ca^2+^ throughout the wing disc pouch decreases during development suggesting an inverse relationship between calcium activity and organ growth rates (Brodskiy et al., 2019). Here, we investigated the role of tissue size in altering Ca^2+^ dynamics to propose an explanation for this finding. To do so, we simulated Ca^2+^ signaling in different sized wing discs. We hypothesized two different possible scenarios that could explain a decrease in integrated Ca^2+^ activity. In the first scenario, the total fraction of initiator cells was allowed to decrease in a power law fashion as development proceeds resulting in a decay in total calcium signaling activity with a transition to predominantly single-cell spiking activity. In the second scenario, gap junction permeability was set to decrease with increasing organ size. Simulations corresponding to each scenario shows an orderly decrease in progression of Ca^2+^ activity starting from ICWs and intercellular transients in smaller simulated discs and decaying to intermittent single-cell spikes in larger simulated discs (Figure 4A, B). This suggests that both scenarios provide a possible explanation for the decrease in integrated Ca^2+^ activity observed in wing discs as development progresses.

**Figure 4:**
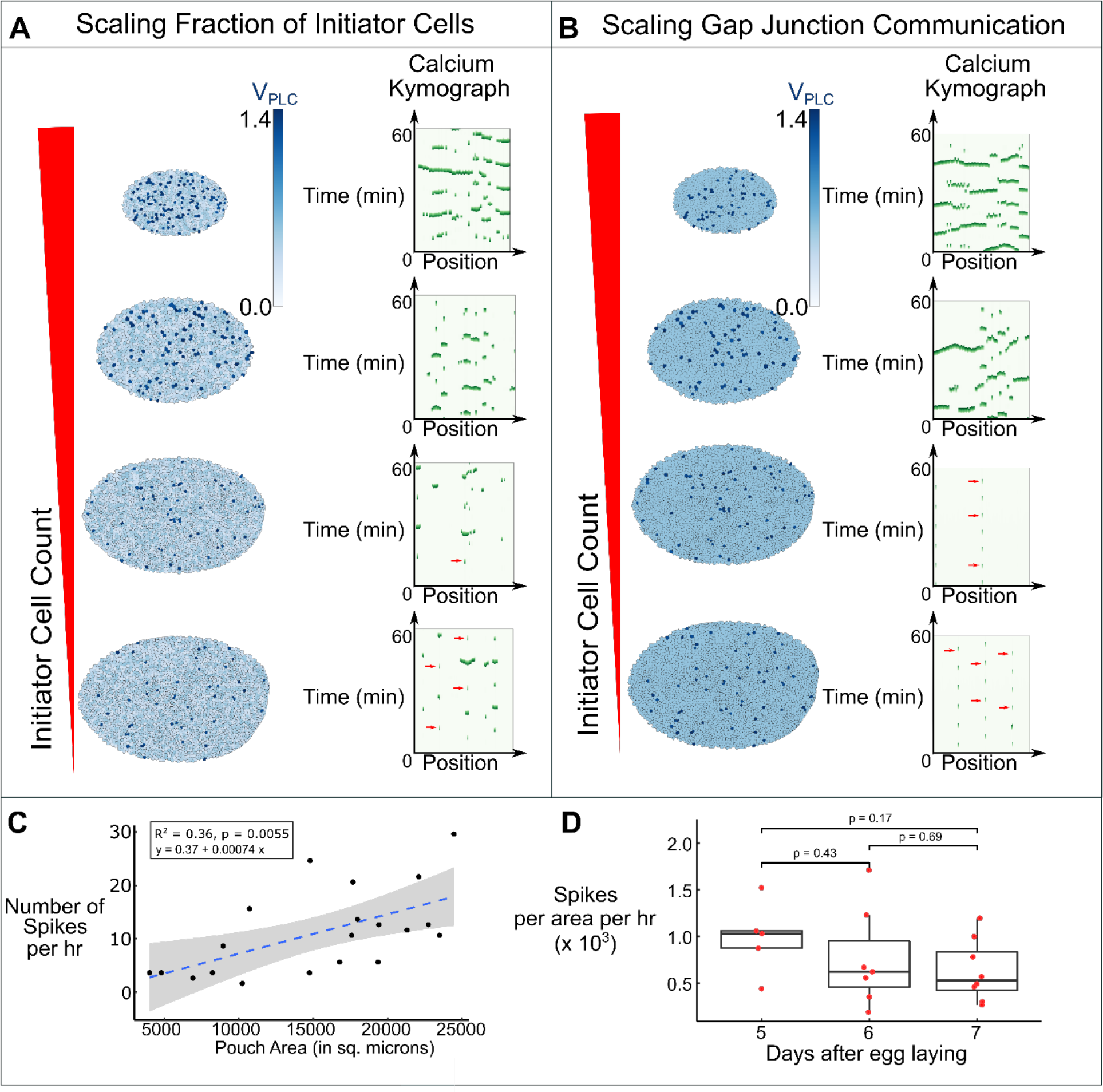
Gap junctional permeability determines total tissue-level signaling activity as development progresses. **(A-B)** Simulations of Ca^2+^ signaling for wing discs of increasing size. In **A**, (*Left column*), the fraction of initiator cells was varied according to the following equation: 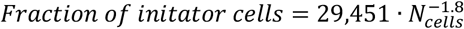. The V_PLC_ range of standby cells was restricted to range from 0.25 – 0.6 with initiator cells being denoted by dark blue cells with V_PLC_ values between 1.3 and 1.5. (*Right column*) Associated 2D kymographs of the simulated pouches shown in **A**. In **B** (*Left column*), the gap junction permeability is varied holding the fraction of initiator cell constant. Gap junction permeability was varied according to the following equation: 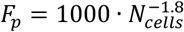 and *F_c_* = 0.1 · *F_p_*. The total number of initiator cells were held constant in this scenario (*N_initiator_* = 65). (*Right column*) Associated kymographs of the simulated pouches. **(C-D)** Experimental validation of the computational predictions in which the discs were cultured ex vivo in Grace’s low 20E media (basal media) for 1 h in the absence of any stimulus triggering Ca^2+^. **(C)** Quantification of total number of Ca^2+^ spikes in different sized wing disc pouch during 1 h culture. A linear regression line was fit to the data set and the p-value for the slope of the fitted line is shown. Since the *p*-value is less than 0.05 (level of significance), a positive correlation between size and the number of spikes could be inferred. Grey region corresponds to 95% confidence bands of the trend line. **(D)** Quantification of total number of spikes in disc pouch scaled with respect to pouch size during various stages of larval development. *p*-values were obtained by Mann Whitney U test.

**Figure 5:**
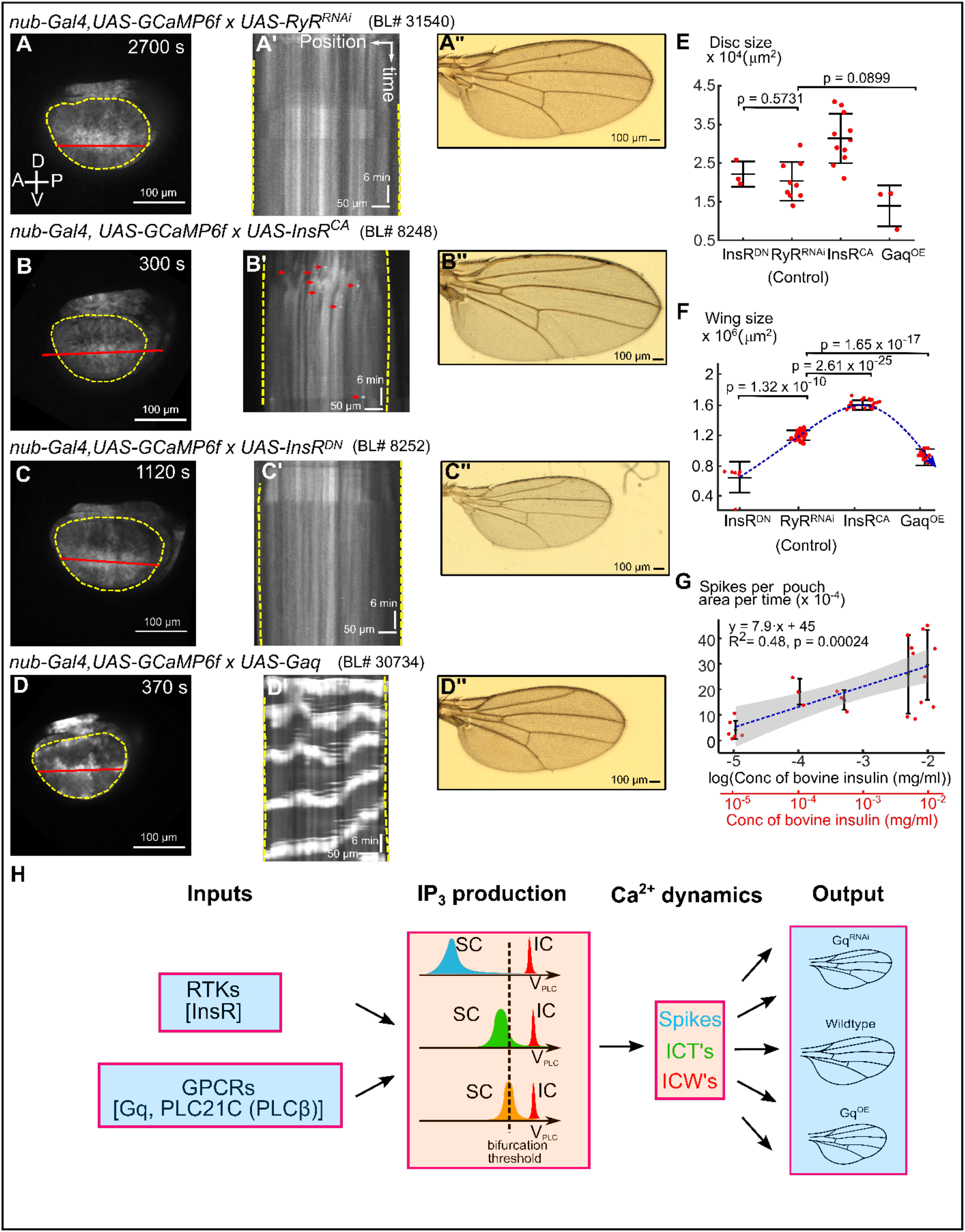
GPCR and insulin signaling regulate wing size and differentially regulate Ca^2+^ signaling. **(A-D)** Montages of time-lapse movies of wing discs cultured ex vivo. **(A’-D’)** Kymographs of the corresponding time-lapse movies. **(A”-D”)** Adult wings from the indicated genetic perturbation. **(E)** Quantification of the wing disc sizes for the indicated genetic perturbations. **(F)** Quantification of the adult wing sizes for the indicated perturbations. **(G)** Quantification of Ca^2+^ spikes when insulin dose is progressively increased in ex vivo cultures. A linear regression trend line was fit to the data and the *p*-value of the slope is shown. Since the *p*-value is less than 0.05, a positive correlation between spikes and the log(concentration) can be inferred. Grey region illustrates 95% confidence bands of the linear regression **(I)** Summary of key findings based on the proposed model for tissue-level regulation of dynamics in epithelial tissues. Scale bars in (A-D) and (A”-D”) represent 100 μm, while yellow dotted lines indicate pouch boundaries, and red lines indicate x-y positions in the kymograph. Horizontal scale bars in (A’-D’) represent 50 μm. Vertical scale bars in (A’-D’) represent 6 min. Student t-test was performed. Labels in (E) represent crosses of UAS-transgene to parental nub>GCaMP6f in the case of InsR^CA^ and InsR^DN^ or nub>GCaMP6f; mcherry in the case of RyR^RNAi^ or Gαq. The UAS lines used are UAS-RyR^RNAi^ (BL#31540), UAS InsR^CA^ (BL#8248), UAS-InsR^DN^ (BL#8252) and UAS-Gq (BL#30734) respectively.

To distinguish between these scenarios, we cultured wing discs from multiple developmental stages from days 5-7 after egg laying (AEL) without any agonist stimulation and observed the resulting Ca^2+^ dynamics. We observed single-cell Ca^2+^ spikes that we interpret as characteristic of “initiator” cells in all samples independent of sizes. We did not observe Ca^2+^ transients or waves in unstimulated smaller wing discs obtained day 5 AEL. We next characterized the total number of spikes in the discs of all sizes. We found a positive correlation between the total number of spikes and the size of the disc pouch (Figure 4C). The difference in spiking activity between discs of varying ages was not significant when scaled for pouch size (Figure 4D). This is consistent with a scenario of a relatively constant number of initiator cells in the system with overall gap junction permeability decreasing as the organ reaches its terminal size. This scenario is further supported by findings from previous reports showing a decrease in gap junctional permeability as larval development proceeds (Weir and Lo, 1982, 1985). Furthermore, a decrease in gap junction permeability increases the net cytosolic residence time and effective concentrations of IP_3_ and Ca^2+^ within the cytosol leading to an increased instance of cytosolic Ca^2+^ spikes.

### Gαq overexpression induces intercellular Ca^2+^ waves and reduces wing size

Next, using the GAL4/UAS system, we overexpressed the wild type splice 3 variant of *Drosophila* Gαq in the wing disc to characterize how different classes of upstream signals impact the spatiotemporal patterns of Ca^2+^ signaling (Ratnaparkhi et al., 2002). G protein-coupled receptor (GPCR) activation stimulates Gαq-driven PLC*β* activity to generate IP_3_ and Ca^2+^ (Hanlon and Andrew, 2015). Strikingly, ectopic Gαq expression was sufficient to generate robust formation of intercellular waves independent of the presence of FEX in the media (Figure 4D, D’). The waves were periodic in nature and were similar to FEX-induced waves (Figure 1C). This experimental finding most likely resembles our previous simulation of ICWs with a small fraction of randomly located initiator cells surrounded by standby cells (Figure 2C).

Strikingly, the wing disc size (day 6 AEL) and adult wing size were significantly reduced when Gαq was overexpressed (Figure 4E-F). To understand whether the reduction in the wing size was due to changes in proliferation or cell growth, we quantified the total number of cells in the region bounded by the LIII, LIV, ACV wing veins and the wing margin. We observed a reduction in the total number of trichomes, which each individual trichome correspond to a cell (García-Bellido et al., 1994). Furthermore, we found that cell size was reduced when Gαq was overexpressed (Figure S5F). However, the wing shape is not significantly affected when Gαq is overexpressed (Figure S5G). In sum, increasing the concentration of Gαq in the pouch is sufficient to generate periodic Ca^2+^ waves. Further, these periodic Ca^2+^ waves are correlated with reduction in wing and disc size suggesting that tissue-wide Ca^2+^ wave activity may play a role in determining final organ size via growth inhibition.

### Insulin signaling increases wing size but only generates localized Ca^2+^ signals

Because FEX is an undefined cocktail of biochemical factors, we tested whether specific ligands added to the organ culture affects Ca^2+^ activity and growth. In addition to FEX, insulin is often added to organ culture media to stimulate cell proliferation (Wyss, 1982; Zartman et al., 2013). Hence, we tested whether insulin signaling regulates Ca^2+^ activity independent of FEX. Similar to other experiments, we upregulated and downregulated insulin signaling in the wing disc using the GAL4/UAS expression system. As expected, wing disc size and adult wing size were decreased when insulin signaling is downregulated (Figure 4B, E, F). Strikingly, we observed that activation of insulin stimulated pathways results in localized Ca^2+^ spikes and ICTs (Figure 4B, B’, C, C’). Titrated the concentration of insulin in the culture media demonstrated that a higher concentration of insulin increased the number of spikes (Figure 4G). Quantification of the Ca^2+^ spikes showed a positive correlation between spikes normalized to area of the pouch with log of insulin concentration (Fig 4G). However, even high concentrations of insulin were not sufficient to generate periodic ICWs. In contrast, expressing a dominant negative form of the insulin receptor resulted in minimal Ca^2+^ spiking activity (Figure 4C, C’) (Hevia et al., 2017). These results demonstrate that insulin signaling stimulates Ca^2+^ activity in the wing disc.

In silico simulations with parameter variations in half activation of the PLC parameter, K_PLC_,and GJ permeability were performed to test scenarios of how insulin signaling can only generate Ca^2+^ spiking activity even under saturating conditions (Figure S3A, S3D, and S6). Although K_PLC_ variations influenced signal frequency, there was no instance in which varying K_PLC_ resulted in production of Ca^2+^ spikes. Single-cell spikes were only observed when gap junction communication was inhibited either experimentally (Figure 3B, C) or in silico (Figure 3A, Figure S3A). Thus, the computational model suggests that insulin signaling must not only stimulate IP_3_ production, but also inhibits gap junction permeability.

As a first step to assess whether Ca^2+^ levels in the cell directly control organ sizes, we exploited the Ca^2+^ buffering effects induced by high levels of GCaMP6f sensor expression. The GCaMP6f sensor consists of a M13 fragment of myosin light chain kinase, GFP and Calmodulin (CaM) to which Ca^2+^ binds (Nakai et al., 2001). To elucidate the role of Ca^2+^ signaling in insulin mediated growth, we compared the effects of co-expressing the GCaMP6f sensor (*K_d_* = 375 ± 14 *nM*) to decrease the amount of free cytosolic Ca^2+^ (Chen et al., 2013). We found that co-expressing GCaMP6f sensor increased wing size when insulin signaling was also stimulated (Figure S7). This increase in size was also observed in control wing disc without insulin upregulation (Figure S7A, B). This analysis provides additional evidence that buffering cytosolic Ca^2+^ signaling influences the final tissue size. In contrast, when we expressed Gαq and GCaMP6f, we did not observe a significant decrease in size when compared to just expressed Gαq without GCaMP6f (Figure S7 C, D). This may be due to the inability of GCaMP6f expression to buffer the high levels of Ca^2+^ in the cytosol when Gαq is overexpressed. We also observed a severe reduction in the wing size, along with vein defects, in flies that were homozygous for the GCaMP6f sensor, consistent with a buffering role for high concentrations of GCaMP6f expression (Figure S8). To validate this finding, we compared the adult wing sizes from flies expressed one and two copies of GCaMP6f. Strikingly, increasing the dose of the relative GCaM6f expression decreased the size (Figure S8 C, D). We observed a severe reduction in size as expression of the transgene was further increased with the presence of two copies of the GAL4 driver. To further elucidate the sponging effects of Ca^2+^, we expressed CaMKII, which binds to Ca^2+^/CaM complex (Yamauchi, 2005). Similar to overexpressing GCaM6f, we observe a significant increase in the wing area (Figure S8E, F). Collectively, these results support a role of cytosolic Ca^2+^ concentration as either a growth enhancer or suppressor and that an optimal amount of Ca^2+^ signaling is required for robust size control.

## Discussion

The main finding of this work is the discovery of a parsimonious mechanistic model that links global hormonal stimulation of Ca^2+^ signaling to emergent spatiotemporal classes of signaling dynamics. Further, we identify downstream correlations to the final organ size that suggest these signaling dynamics may mediate growth-related information used by the system to tune organ size. To do so, we developed a geometrically accurate computational model of Ca^2+^ signaling in the *Drosophila* wing imaginal disc, a premier model for studying conserved cell signaling mechanisms (Hariharan, 2015; Johnston and Gallant, 2002; Potter, 2001). Previously, we discovered a correlation between Ca^2+^ signaling during larval growth and final organ size. Four distinct patterns of Ca^2+^ signaling activity occur in the wing imaginal disc pouch as observed in vivo and ex vivo (Brodskiy et al., 2019). Through systems-level computational analysis, we established the essential conditions required for generating the different patterns.

The model predicts two cell types with different levels of IP_3_ production: “initiator cells” with high IP_3_ production and “standby cells” with baseline IP_3_ production levels. The presence of initiator cells is necessary, but not sufficient, to induce multicellular Ca^2+^ signaling. Additionally, the distribution of the maximal rate of IP_3_ production, V_PLC_, in the standby cells determines the spatial range of Ca^2+^ signaling. As V_PLC_ values of standby cells approaches the Hopf bifurcation threshold, Ca^2+^ activity transitions from single-cell signals toward global signals.

What are the possible functional implications of a tissue consisting of initiator and standby cells? A recent study on electrical signal transmission in bacterial communities suggests that the transition from localized short-range signaling to global community-level communication is associated with a cost-benefit balance (Larkin et al., 2018). In that context, long-range signaling increases the overall fitness of the community against chemical attacks, while a reduction in growth rate is the cost to individual cells. In the wing disc, a similar analogy can be drawn where significant generation of long-range Ca^2+^ signals due to overexpression of Gαq results in reduced wing disc growth. Thus, the proposed model in this work can also be characterized as a cost-benefit tradeoff within the context of tissue-level signaling. For instance, it has been suggested that the fast Ca^2+^ waves facilitate migration and proliferation of the healing cells by inhibiting excessive apoptotic response during wound healing in epithelia (Justet et al., 2016).

Our model also predicts that the inhibition of GJs lowers the Hopf threshold necessary for generating Ca^2+^ spikes. We have validated this prediction experimentally where inhibition of GJs results in the formation of Ca^2+^ spikes in the absence of external agonist. Further, computational simulations demonstrate that as GJ permeability decreases, there is a transition of activity from synchronous global to asynchronous local Ca^2+^ activity. How gap junctional mediated Ca^2+^ signaling is connected to the regulation of cell mechanics is not currently understood and warrants further investigation. One possibility is that tension impacts the level of gap junction communication between cell and may also influence the activity of mechanosensitive ion channels (Kaouri et al., 2019). Feedback between calcium signaling and cell mechanics may play important roles in ensuring tissue growth and morphogenesis. This is evident from our previous experimental work and other experimental studies in the literature where knockdown of gap junctional proteins such as Inx2 leads to a reduction in wing and eye size in *Drosophila* (Brodskiy et al., 2019; Richard et al., 2017).

Upregulated insulin signaling increases the formation of Ca^2+^ spikes. One possible implication is that insulin signaling inhibits GJs in addition to increasing PLC*γ* activity. This implication is consistent with the role of insulin signaling in inhibiting gap junction proteins Connexin43 (Homma Nobuo et al., 1998; Lin et al., 2003; Warn-Cramer and Lau, 2004). Thus, the increase of adult wing and developing wing disc size from higher insulin activity correlates with higher levels of localized Ca^2+^ spiking activity, which has a limited total integrated calcium signal at that tissue level. In contrast, Ca^2+^ waves induced by Gαq overexpression are correlated with smaller adult wings and wing discs. Higher Gαq expression, which activates PLC*β* activity results in robust production of tissue-scale intercellular Ca^2+^ waves. We show experimentally that insulin signaling controls Ca^2+^ spike activity in the wing disc, potentially through GJ inhibition or PLC*γ* activation, whereas GPCR-based Gαq signaling is sufficient to generate global Ca^2+^ waves. These results are consistent with previous reports of Ca^2+^ spikes being observed in discs that have reached their final size, and Ca^2+^ waves being observed in smaller developing discs (Brodskiy et al., 2019). A recent study reported after we archived this study as a preprint found that Ca^2+^ signals are initiated in response to wounding by the G-protein coupled receptor Methuselah like 10 (Mthl10) (O’Connor et al., 2020). Mthl10 is activated by Growth-blocking peptides (Gbps) (O’Connor et al., 2020). Whether Mthl10 also is involved in developmental growth requires further investigation, but may be consistent with our findings that intercellular calcium wave activity inhibit organ growth. These findings suggest that Ca^2+^ acts as both a growth enhancing and growth inhibiting signal dependent upon the tissue-level scale of the activity and level of gap junction coupling.

The spatial range of tissue-level signaling is determined by how the IP_3_ production is organized with respect to a Hopf bifurcation threshold throughout the tissue. Localized transients are correlated with larger wings induced by insulin-stimulated growth. Global signaling is correlated with smaller wings that are stimulated by upstream GPCRs and Gαq upregulation. This resembles a paradoxical network motif where the Ca^2+^ signaling has two opposite effects on the same target, dependent upon the tissue-level scale of the Ca^2+^ signal. Within the context of the hypothesized IP_3_/shunt model proposed in our previous study, the strong induction of Ca^2+^ waves will reduce the level of phosphatidylinositol 4,5-bisphosphate (PIP_2_), a key substrate for growth (Brodskiy et al., 2019). This may occur as high levels of Gαq/PLC activity are proposed to deplete PIP_2_ levels (Loew, 2007). This is likely due to substrate depletion of PIP_2_ through promotion of IP_3_ generation and downstream activity by stimulation of PLC activity. In turn, this would lead to reduced availability of PIP_2_ for conversion of PIP_2_ to phosphatidylinositol-trisphosphate (PIP_3_), a key second messenger for stimulating protein kinase AKT and downstream growth promotion (Czech, 2000).

Similar to the reduced wing size observed in this study due to overexpression of Gαq, we have also reported that knockdown of Gαq gene decreases wing size in our previous study (Table S3) (Brodskiy et al., 2019). Comparing the reduction in wing size due to perturbation of Ca^2+^ signaling components with known size control genes such as morphogens (Dpp, Wg, Hippo) and mechanical transducers (RoK) indicate that the reduction in size is comparable to when Ca^2+^ signaling is perturbed (Table S3). Taken together, these experimental findings imply that Gαq signaling is paradoxical in nature. In the context of conserved network motif observed in biological systems, paradoxical components have the ability to activate and inhibit the downstream target despite a single source of stimulus (Hart and Alon, 2013). Gαq could possibly be a growth promoter and growth inhibitor. This correlation motivates further studies that map out the exact molecular players that are downstream of Gαq signaling. This will require careful quantification of PIP_2_ and PIP_3_ under genetic perturbations of GPCR signaling. Additionally, future work is needed to quantify the metabolic benefits and costs of Ca^2+^ signaling during tissue growth to observe if abundant use of metabolic resources to consistently propagate global activity is explanatory for the reduction in size in Gαq overexpression wings.

## Materials and Methods

### Drosophila genetics

We used the GAL4/UAS system to express modulators of the Ca^2+^ signaling pathway in the wing disc (Brand and Perrimon, 1993; Duffy, 2002). A nub-GAL4, UAS-GCaMP6f reporter tester line was created by recombining nub-GAL4 and UAS-GCaMP6f lines (Narciso et al., 2015). Additionally, a second tester line was used that also includes UAS-mcherry. Gene perturbations were generated by crossing the tester line to either RNAi-based transgenic lines (UAS-Gene X^RNAi^) or gene overexpression (UAS-Gene X). The following UAS transgenic lines were used: UAS-RyR^RNAi^ (BL#31540) (Perkins et al., 2015), UAS-Gq (BL#30734) (Perkins et al., 2015), UAS-InsR^CA^ (BL#8248) (Werz et al., 2009), UAS-InsR^DN^ (BL#8252) (Kakanj et al., 2016). Progeny wing phenotypes are from F1 male progeny emerging form the nub-Gal4, UAS-GCaMP6f/CyO x UAS-X cross or nub-Gal4, UAS-GCaMP6f/CyO; UAS-mcherry x UAS-X cross. Flies were raised at 25 °C and on a 12-hour light cycle.

### Live imaging

Wandering third instar larva approximately 6 days after egg laying were dissected in ZB media with 15% fly extract to obtain wing discs (Brodskiy et al., 2019). ZB media + 15% fly extract contains 79.4% (v/v) ZB media, 0.6% (v/v) of 1 mg/ml of insulin (Sigma aldrich), 15% ZB-based fly extract and 5% penicillin/streptomycin (Gibco). Wing discs were loaded into the previously described REM-Chip (Narciso et al., 2017) and imaged using Nikon Eclipse Ti confocal microscope with a Yokogawa spinning disc and MicroPoint laser ablation system. Image data were collected on an IXonEM+colled CCD camera (Andor technology, South Windsor, CT) using MetaMorph v7.7.9 software (Molecular devices, Sunnyvale, CA). Discs were imaged at three z-planes with a step size of 10 μm, 20x magnification and 10-seconds intervals for a total period of one hour, with 200 ms exposure time, and 50 nW, 488 nm laser exposure at 44 % laser intensity. We blocked GJ by inhibiting innexin-2 using Carbenoxolone (Cbx, Sigma Aldrich) drug (Narciso et al., 2015). Wing discs were incubated in ZB + 15% FEX with 30 μM Cbx for one hour before imaging. To induce Ca^2+^ transients, we imaged wing discs in ZB media + 2.5 % FEX (Burnette et al., 2014). Ca^2+^ waves were induced by imaging the wing disc in ZB media + 15% FEX. Ca^2+^ fluttering was observed when discs were imaged in ZB media + 40% FEX respectively.

### Image processing

All the images were processing in FIJI. Volume viewer plugin was used to generate 3D Kymographs. Briefly, TIFF stacks had background subtracted using the rolling ball background subtraction algorithm with a rolling ball radius of 15. The TIFF stack was then processed by the volume viewer plugin. Stacks were adjusted using the base Fire LUT setting to portray the signaling intensities. A similar approach was followed for the simulation outputs.

### Model formulation

Figure 1 summarizes the experimental system and data. Different classes of patterns emerge at the tissue-level as the level of global stimulation increases: spikes, intercellular Ca^2+^ transients (ICTs), intercellular Ca^2+^ waves (ICWs) and global fluttering (Brodskiy et al., 2019). However, a mechanistic understanding linking hormonal stimulation levels to transitions in these qualitative classes of organ-level signaling is lacking. We therefore formulated a computational model to test mechanistic hypotheses that could explain the observed Ca^2+^ signaling dynamics.

### Intracellular model

A modified model of Ca^2+^ signaling toolkit is based on adaptation of previous single-cell model of calcium signaling (Politi et al., 2006). The model is summarized in Figure 1E. A more comprehensive description can be found in Supporting Information. To recapitulate the same time resolution as the experiments, the simulation time is 1 hour and for generating videos, samples are obtained every 10 s.

### Tissue model

For constructing a realistic model of the tissue, we used experimental images of a wing pouch to build an accurate model of the tissue structure. Figure S1 depicts the structure of the tissue used for simulations and the statistics of the corresponding network. More details on the geometry of the model are discussed in Supporting Information.

### Multicellular model

The realistic model of the tissue was combined with proposed intracellular model. Diffusion of the second messengers IP_3_ and Ca^2+^ between adjacent cell was incorporated into the 2D model. Therefore, the tissue level model is a system of coupled ODE’s. A complete description of the model is provided in Supporting Information.

### Quantification and Statistical Analysis

#### Quantification of adult wings and statistics

Total wing area was measured using ImageJ. We traced the wing margin by following veins L1 and L5 and the wing hinge region was excluded from the size analysis. All statistical analyses were performed using MATLAB or R. For comparisons, we used student t-tests to assess the statistical significance. *p*-value, standard deviation and sample size (n) are as indicated in each figure and legend.

## Data and Code Availability

All the data and simulation codes are available on our lab GitHub repository: https://multicellularsystemslab.github.io/MSELab_Calcium_Cartography_2021/

## Acknowledgements

The work in this paper was supported by NIH Grant R35GM124935 and NSF Award CBET-1553826. The authors gratefully acknowledge the Notre Dame Center for Research Computing (CRC) for providing computational facilities. The authors would also like to thank members of the Zartman lab for helpful discussions.

