## Supplementary material for "From spikes to intercellular waves: tuning intercellular Ca^2+^ signaling dynamics modulates organ size control": SI text

**Dharsan K. Soundarrajan[1,†], Francisco J. Huizar[1,2, †], Ramezan Paravitorghabeh[1,†], Trent Robinett[1], Jeremiah J. Zartman[1,2*]**
[1]Department of Chemical and Biomolecular Engineering, University of Notre Dame, South Bend, Indiana
[2]Bioengineering Graduate Program, University of Notre Dame, South Bend, Indiana

[†]These authors contributed equally

### Computational model

Several mathematical models have been proposed to describe intra- and intercellular $Ca^{2+}$ wave propagation (Sera and Kudo, 2020). The base computational model extends a previous formulated model reported by Politi and colleagues to describe single-cell $Ca^{2+}$ oscillations observed in Chinese hamster ovarian (CHO) cells (Politi et al., 2006). This model accounts for the formation and degradation of $IP_3$, $Ca^{2+}$ flux across the endoplasmic reticulum (ER) through $IP_3R$, and sarco/endoplasmic reticulum $Ca^{2+}$-ATPase (SERCA), $IP_3R$ and ER $Ca^{2+}$ dynamics. The model consists of four state variables: cytosolic $IP_3$ ($p$), cytosolic $Ca^{2+}$ ($c$), the ER $Ca^{2+}$ concentration ($s$), and the fraction of $IP_3$ receptors that have not been inactivated by $Ca^{2+}$ ($r$).

#### *$IP_3$ dynamics*

$IP_3$ is generated in the cytosol by phospholipases (PLC) (Berridge and Irvine, 1989). The *Drosophila* genome consists of three PLC genes. They include *PLC21C* and *norpA*, which are related to the PLCβ1-4 subfamily of the *Homo sapien* homologs, and a single PLCγ (*sl*) (Balakrishnan et al., 2015). While different classes of PLC can hydrolyze $PI(4,5)P_2$ to generate $IP_3$ and DAG, they are activated by different receptors on the cell surface. For instance, PLC21C is activated by the heterotrimeric G-protein αq subunit in response to G-protein receptor signaling. On the other hand, PLCγ is recruited via its SH2 domain to activated receptor tyrosine kinase at the plasma membrane (Balakrishnan et al., 2015). In our model, we describe all the combined production of $IP_3$ as dependent on the total enzymatic activity of the cell:

$$v_{PLC} = V_{PLC} \frac{c^2}{K_{PLC}^2 + c^2} \tag{1}$$

where $V_{PLC}$ describes the maximal production rate of $IP_3$ and $K_{PLC}$ describes the sensitivity of PLC to $Ca^{2+}$. Furthermore, it is essential to note that the parameter $V_{PLC}$ depends on agonist concentration, which is directly proportional to PLC substrate $PIP_2$, and we assume that $V_{PLC}$ describes a net activity of PLC activated by upstream receptor. While different PLC homologs are structurally different and are not activated by the same receptors upstream, our model assumes that both PLCs are described by the above equation with same parameters. In many cell types, $IP_3$ production rate depends on both the concentration of PLC and $Ca^{2+}$. Since cytosolic $Ca^{2+}$ is both a positive and negative regulator of PLC activity, we assume only consider the case of positive regulation of $IP_3$ production by $Ca^{2+}$.

Our model also considers degradation of $IP_3$ by other factors such as $IP_3$ kinases, which converts $IP_3$ to $IP_4$. We generalize the degradation of $IP_3$ using first order kinetics. Collectively, the equation describing the dynamics of $IP_3$ is:

$$\frac{dp}{dt} = J_p + v_{PLC} - k_{5P}p \tag{2}$$

where $J_p$ is the flux of $IP_3$ through gap junctional communication. We assume that $IP_3$ diffuses from one cell to the adjacent cells through gap junctional coupling. We model the flux through gap junctions (GJs) using the following equation

$$J_p \approx F_p\left[\sum\nolimits_{j\in N_i} p_j l_j - p_j\left(\sum\nolimits_{j\in N_j} l_j\right)\right] \tag{3}$$

where $F_p$ refers to the permeability of $IP_3$ through gap junctional communication, $p_j$ refers to $IP_3$ concentration in neighboring cell $j$ and $l_j$ refers to the length of cell boundary shared by cells $i$ and $j$ respectively. We assume that the intracellular diffusion of $IP_3$ is fast relative to the diffusion of $IP_3$ between cells through GJs. Consequently, we have neglected terms that describe intracellular diffusion.

*$Ca^{2+}$ dynamics*

$Ca^{2+}$ is released through $IP_3R$ from the ER. Similarly, cytosolic $Ca^{2+}$ is pumped into the cytosol using SERCA pumps. In many cell types, $Ca^{2+}$ is also pumped out from cytosol to extracellular space through plasma membrane. In our model, we ignore the flux of $Ca^{2+}$ through the cell's plasma membrane, and we only consider the transport of $Ca^{2+}$ from ER to cytosol. To describe the $IP_3R$ dynamics, we use the form of equation used by Li and Rinzel (Li and Rinzel, 1994). Thus, the dynamics of cytosolic $Ca^{2+}$ is given by

$$\frac{dc}{dt} = J_c + [k_1\left(r.\frac{c}{K_a + c}\frac{p}{K_p + p}\right)^3 + k_2](s - c) - V_{SERCA}\frac{c^2}{c^2 + K_{SERCA}^2} \tag{4}$$

where $k_1$ refers to maximal rate of $Ca^{2+}$ release, $K_a$ is the rate constant characterizing $Ca^{2+}$ binding to activating site in $IP_3R$, $K_p$ is the rate constant characterizing $IP_3$ binding to $IP_3R$, $k_2$ refers to $Ca^{2+}$ leak out of ER, $V_{SERCA}$ is the maximum rate of SERCA pump and $K_{SERCA}$ is the half activation constant. We assume that $Ca^{2+}$ acts as both a positive and negative regulator of $IP_3R$ which is consistent with experimental observations of single channel properties of wild type *Drosophila* receptor that has been studied using lipid bilayer reconstitution technique (Srikanth et al., 2004). Similar to $IP_3$, we also model diffusion of $Ca^{2+}$ through GJs by the following equation

$$J_c \approx F_c \left[ \sum\nolimits_{j \in N_i} c_j l_j - c_j \left( \sum\nolimits_{j \in N_j} l_j \right) \right] \quad (5)$$

where $F_c$ refers to the permeability of $Ca^{2+}$ through GJs, $m_j$ refers to the concentration of $Ca^{2+}$ in neighboring cell $j$ and $l_j$ refers to the length of cell boundary shared by cells $i$ and $j$ respectively.

Similarly, we describe the dynamics of $Ca^{2+}$ concentration in the ER as

$$s(t) = \frac{c_{tot} - c(t)}{\beta} \quad (6)$$

where $s$ is the $Ca^{2+}$ concentration in the ER and $\beta$ is the ratio of effective cytoplasmic and effective ER volume, $c$ refers to the cytosolic $Ca^{2+}$ concentration in the cell and $c_{tot}$ refers to the total $Ca^{2+}$ concentration in the cell which includes both ER and the cytosol.

*$IP_3R$ dynamics*

We assume that the $Ca^{2+}$ binding to the inactivating site on the $IP_3R$ is a slow process. Consequently, we consider the dynamics of $IP_3R$ inactivation by $Ca^{2+}$ as a separate differential equation given below

$$\tau_r \frac{dr}{dt} = \left[ 1 - r \frac{(K_i + c)}{K_i} \right] \quad (7)$$

where $r$ refers to the fraction of $IP_3R$ that is not inactivated by $Ca^{2+}$, $K_i$ refers to the binding coefficient characterizing $Ca^{2+}$ binding to the inactive site on the $IP_3R$ and $\tau_r$ refers to the characteristic time of $IP_3R$ inactivation.

**Index of Supplemental figures:**

Figure S1: Computational framework

Figure S2: Single-cell $Ca^{2+}$ dynamics

Figure S3: Intercellular $Ca^{2+}$ communication is altered by changing $F_p, F_c, K_\tau, \tau_{max}, K_{PLC}$.

Figure S4: Bifurcation analysis of the modified model

Figure S5: Overexpression of Gαq decreases cell number and cell size

Figure S6: GJ inhibition is the key driver of single-cell $Ca^{2+}$ spike activity

Figure S7: Sponging cytosolic $Ca^{2+}$ increases overall wing size

Figure S8: Increasing gene dosage of $Ca^{2+}$ sensor GCaMP6f dramatically reduces wing size

Table S1: Baseline parameters used in the model

Table S2: Extended data movies

Table S3: Changes in wing area for known perturbations through Gal4/UAS system. Note that maximal deviations wing size for strong growth perturbations is in range of 20-50%.

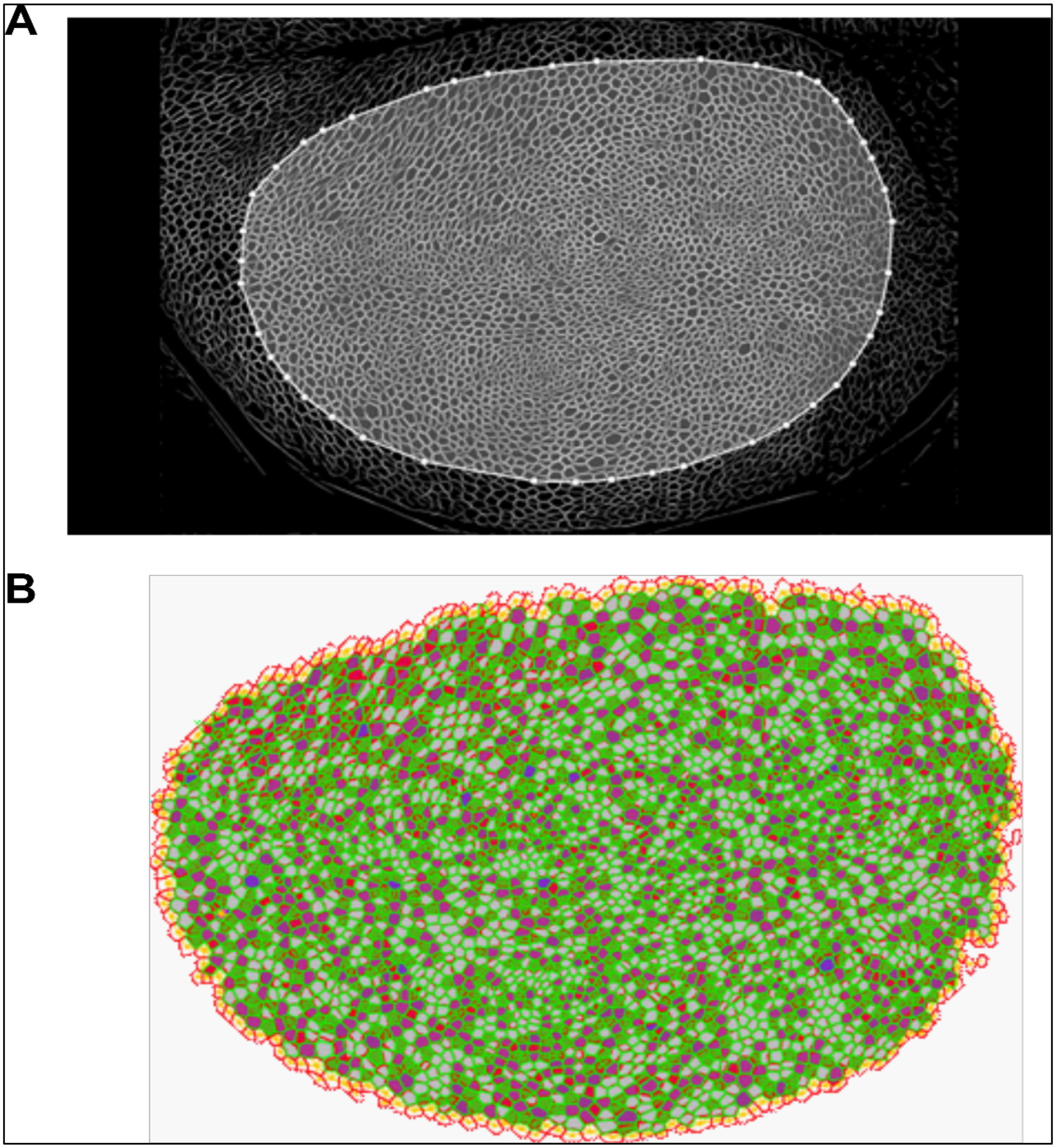


**Figure S1: Computational framework. (A)** Experimental *Drosophila* imaginal disc showing cell boundaries marked with Ecad::GFP. The developing wing pouch has been segmented using ImageJ. The genotype of the *Drosophila* used is *yw;;dECad::GFP* (BL# 46556) **(B)** A pouch constructed computationally using EpiTools that served as a basis for $Ca^{2+}$ signaling simulations. In brief, cells were segmented from a wing disc. Centroids of segmented cells were used to define cellular positions in the simulated wing disc. A Voronoi tessellation followed by multiple rounds of Lloyd's relaxation (Lloyd, 1982) was used to define a template wing disc that matches the experimentally observed network topology.

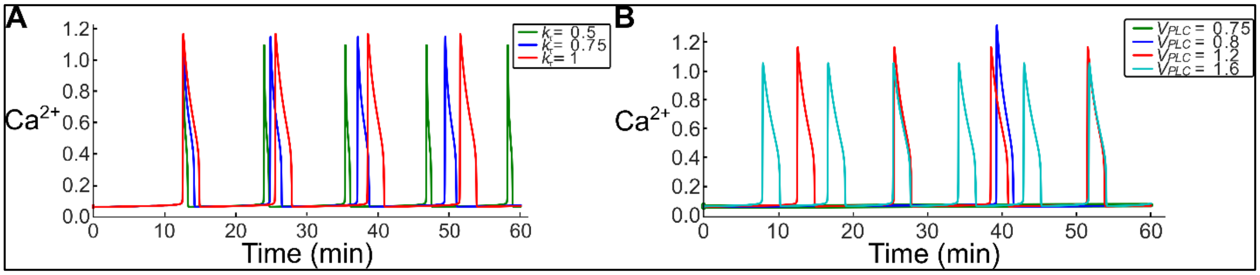


**Figure S2: Single-cell $Ca^{2+}$ dynamics**. The model was calibrated to match experimental single-cell frequency and amplitude. Perturbations to $k_\tau$ **(A)** alters the frequency of $Ca^{2+}$ oscillations whereas stimulation strength $V_{PLC}$ **(B)** alters the frequency and amplitude, respectively.

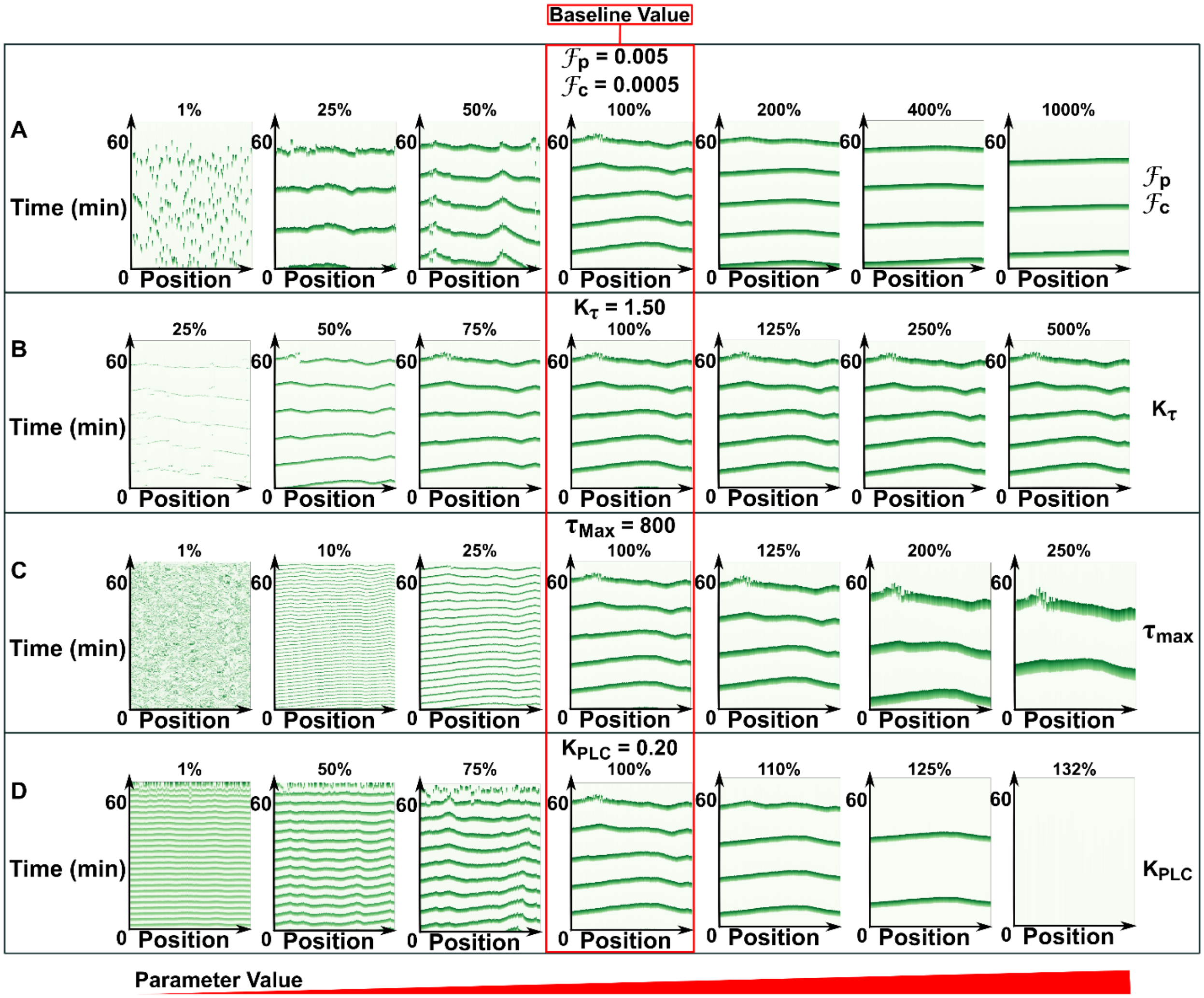


**Figure S3: Intercellular $Ca^{2+}$ communication is altered by changing** $F_p, F_c, K_\tau, \tau_{max}, K_{PLC}$. Four different parameters in the 2D model were varied. The baseline parameters for the computational model were selected to generate intercellular waves. This parameter set was selected to compare how parameter variations changed the signal dynamics: $IP_3$ GJ permeability ($F_p$) of 0.005 $\mu M^2 s^{-1}$, $Ca^{2+}$ GJ permeability ($F_c$) of 0.0005 $\mu M^2 s^{-1}$, $k_\tau$ of 1.50 µM, $\tau_{max}$ of 800 $s^{-1}$, and $K_{PLC}$ of 0.20 µM (red box). Simulations were performed varying only one parameter value while holding all others constant. **(A)** GJ permeability of $IP_3$ and $Ca^{2+}$ influences synchronization of $Ca^{2+}$ signaling among cells. Decrease from baseline results in a transition of intercellular waves to intercellular transients, and to single-cell spikes. **(B, C)** Varying the $k_\tau$ and $\tau_{max}$ parameters influenced the characteristic time associated with inactivation of $IP_3$. Decrease in one or the other results in a decrease of the WHM of $Ca^{2+}$ signaling transients in cells. Only an increase in $\tau_{max}$ resulted in a decrease of frequency and increase in WHM. **(D)** Variations in the half-activation of $V_{PLC}$ term ($K_{PLC}$) only changed the frequency of the ICWs.

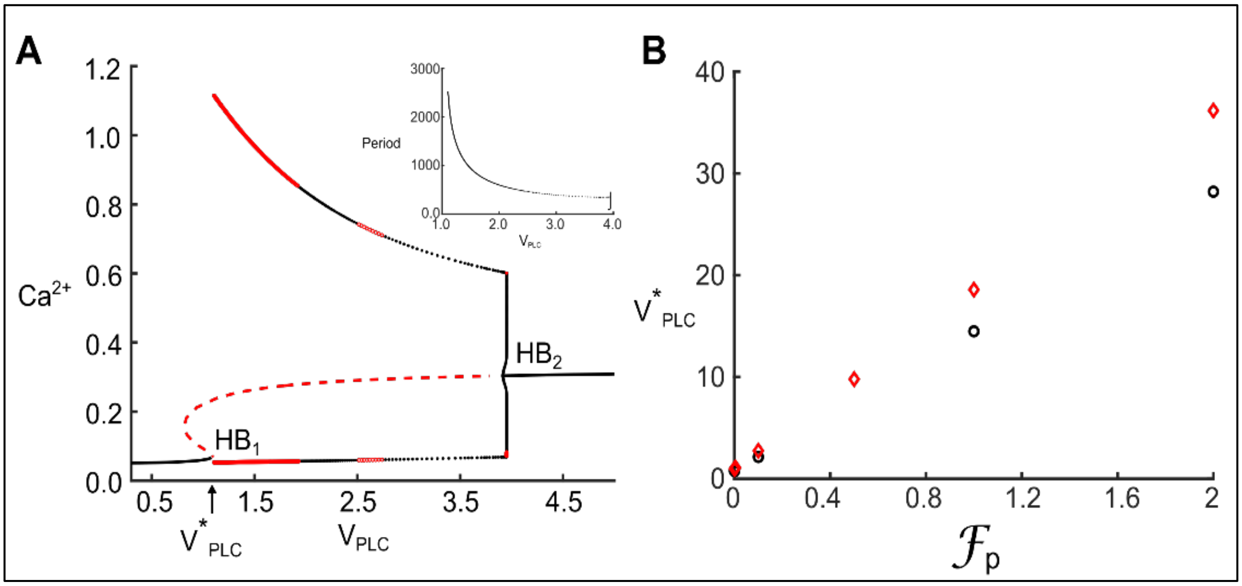


**Figure S4: Bifurcation analysis of the modified model (A)** Bifurcation diagram for the modified model used in this study; shown are the maxima and minima of the $Ca^{2+}$ oscillations (dots) and the $Ca^{2+}$ steady states (solid and dashed lines) as a function of the stimulus ($V_{PLC}$). Solid and dashed lines in red indicate stable and unstable states, respectively. Red dots indicate the maxima and minima of unstable limit cycle and the black dots indicate maxima and the minima of the stable limit cycle. HB, Hopf bifurcation occurs when $V_{PLC}$ is varied. Inset figure shows the period of $Ca^{2+}$ oscillations as a function of $V_{PLC}$. **(B)** Blocking permeability of $IP_3$, $F_p$ via gap junctions decreases $V_{PLC}$ where the initial Hopf bifurcation point ($HB_1$) occurs in the bifurcation diagram. Block dots indicate conditions where permeability of $Ca^{2+}$, $F_c$ is set to 0. Red diamonds indicate conditions where $F_c$ is set to 0.5.

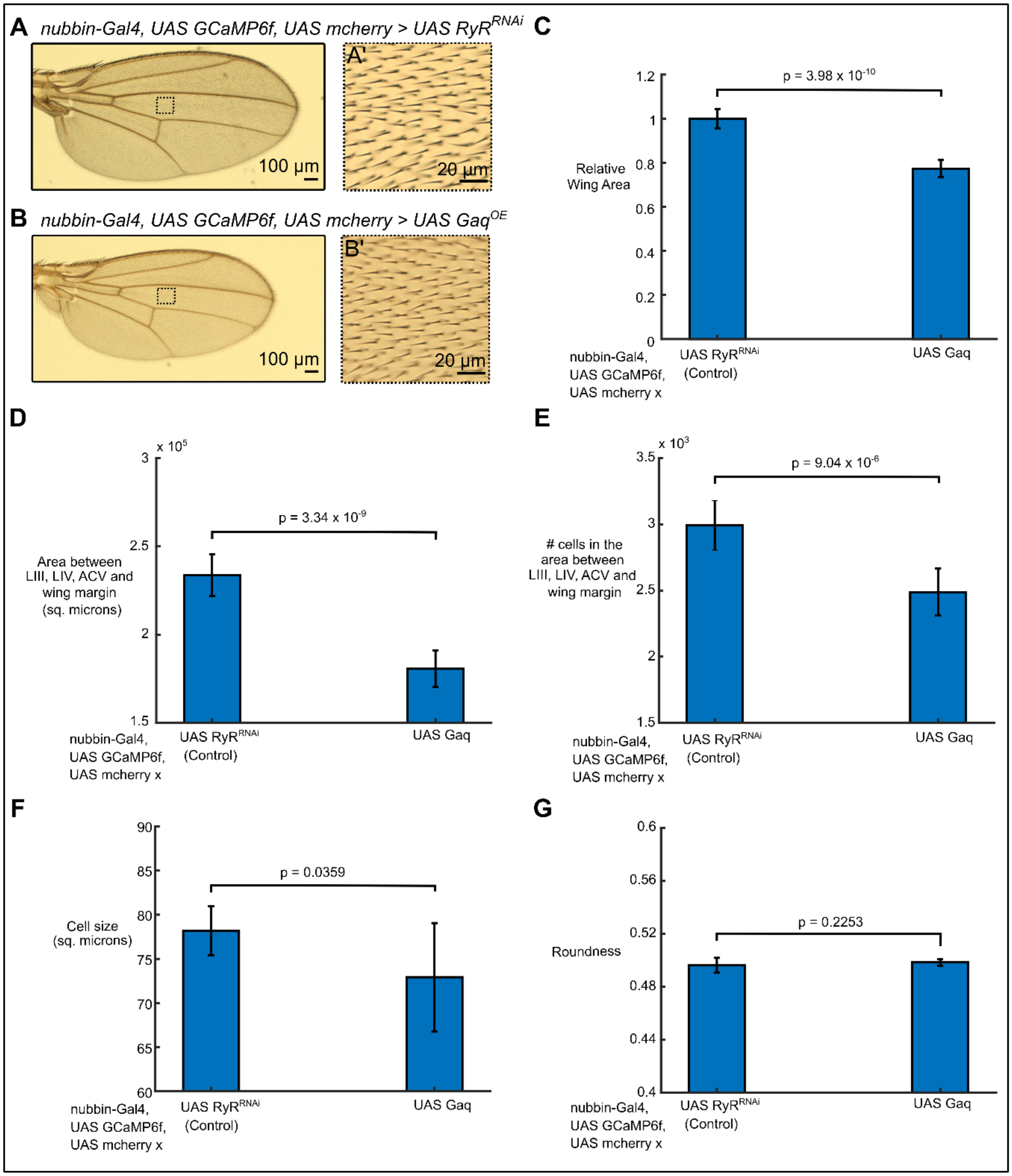


**Figure S5: Overexpression of Gαq decreases cell number and cell size. (A-B)** Wings from adult males expressing *RyR*$^{RNAi}$ and wild type Gαq splice 3 variant with *nubbin-Gal4*, *UAS GCaMP6f*, *UAS mcherry*. **(A'-B')** Region of interest (ROI) where the total number of setae was calculated. **(C-F)** Quantification of the wing size defined here as the area bounded by LIII, LIV, ACV and the wing margin, total cell number and cell area. Overexpression of Gαq in the pouch results in a decrease in total wing area, cell number and cell size. 10 samples were analyzed per condition. Error bars represent standard deviation. **(G)** Quantification of roundness of the adult wing. Gαq overexpression does not affect the roundness. Student t-test was used for statistical significance testing.

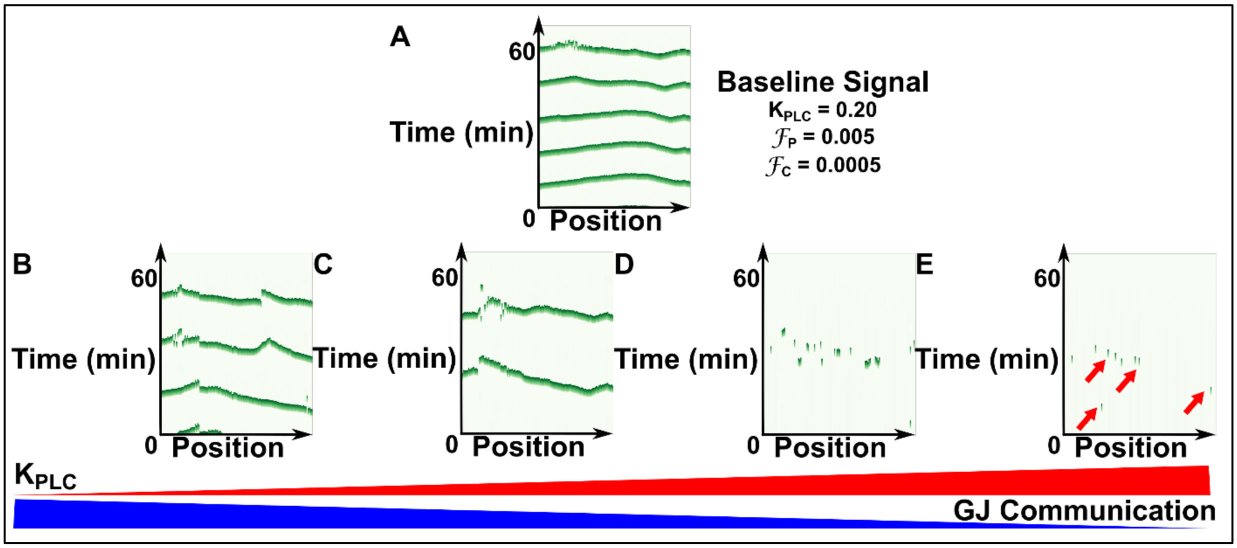


**Figure S6: GJ inhibition is the key driver of single-cell $Ca^{2+}$ spike activity.** To replicate the ex vivo observations of insulin inducing single-cell $Ca^{2+}$ spikes, GJ permeability and the half-activation of $V_{PLC}$ were varied simultaneously in silico. **(A)** A baseline intercellular wave was used as the comparison for how parameter variations changed signal with the following parameter values: $IP_3$ gap junction permeability ($F_p$) of 0.005 μ$M^2s^{-1}$, $Ca^{2+}$ gap junction permeability ($F_c$) of 0.0005 μ$M^2s^{-1}$, and $K_{PLC}$ of 0.20 μM. **(B-E)** $K_{PLC}$ is increased left-to-right (red bar), and gap junction communication is decreased left-to-right (blue bar). An increase in $K_{PLC}$ results in a decrease in frequency, while decrease in gap junction communication results in single-cell $Ca^{2+}$ spikes (red arrows).

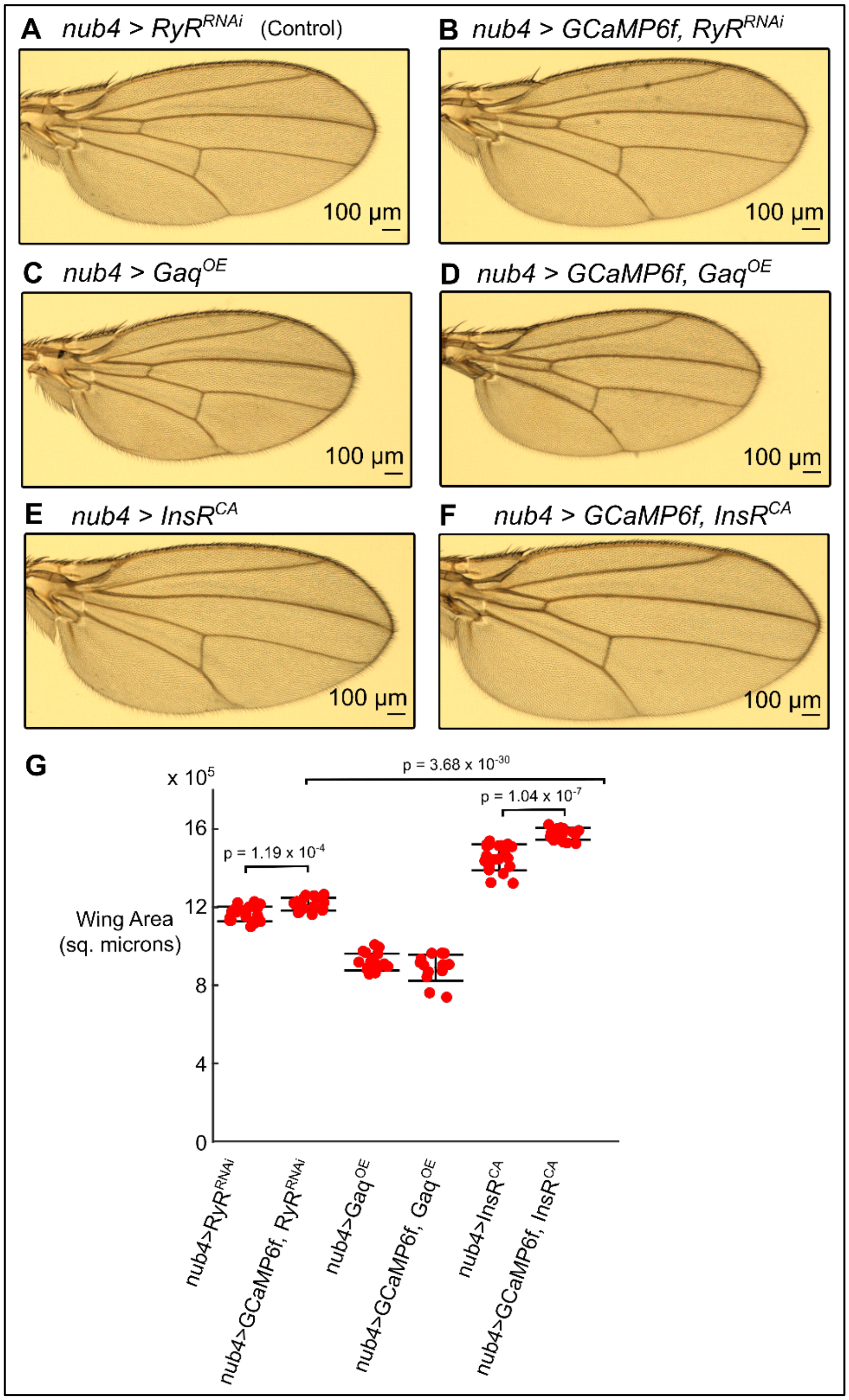


**Figure S7: Sponging cytosolic $Ca^{2+}$ increases overall wing size**. **(A-F)** Wings from adult males with the indicated crosses. **(A)** *nubbin-GAL4* x *UAS-RyR$^{RNAi}$* (i.e., nub4>RyR$^{RNAi}$), **(B)** *nubbin-GAL4, UAS-GCaMP6f* x *UAS-RyR$^{RNAi}$* (i.e., nub4>GCaMP6f,RyR$^{RNAi}$), **(C)** *nubbin-GAL4* x *UAS-Gaq$^{OE}$* embryonic splice 3 variant of Gaq (i.e., nub4>Gaq$^{OE}$), **(D)** *nubbin-GAL4*, *UAS-GCaMP6f x UAS-Gaq$^{OE}$* (i.e., nub4>GCaMP6f,Gaq$^{OE}$), **(E)** *nubbin-GAL4 x UAS-InsR$^{CA}$* (i.e., nub4>InsR$^{CA}$) gain of function mutant where the α subunit is partially deleted **(F)** *nubbin-GAL4*, *UAS-GCaMP6f x InsR$^{CA}$* (i.e., nub4>GCaMP6f, InsR$^{CA}$). **(G)** Quantification of adult wings. The genetic encoded calcium sensor, GCaMP6f, binds to $Ca^{2+}$ with high affinity, thus expression of the sensor will to some degree act as a sponge of cytosolic $Ca^{2+}$. Interestingly, the presence of the GCaMP6f sponge with constitutively activated insulin signaling increases the adult wing size (E,F).

Similar enhancement of wing size was observed in control wings when GCaMP6f sensor was expressed (A,B) No significant change in the adult wing size was observed when Gαq was overexpressed, suggesting sponging effects are trivialized under Gαq overexpression (C,D). Unpaired student t-test was used, and the p-values are indicated above.

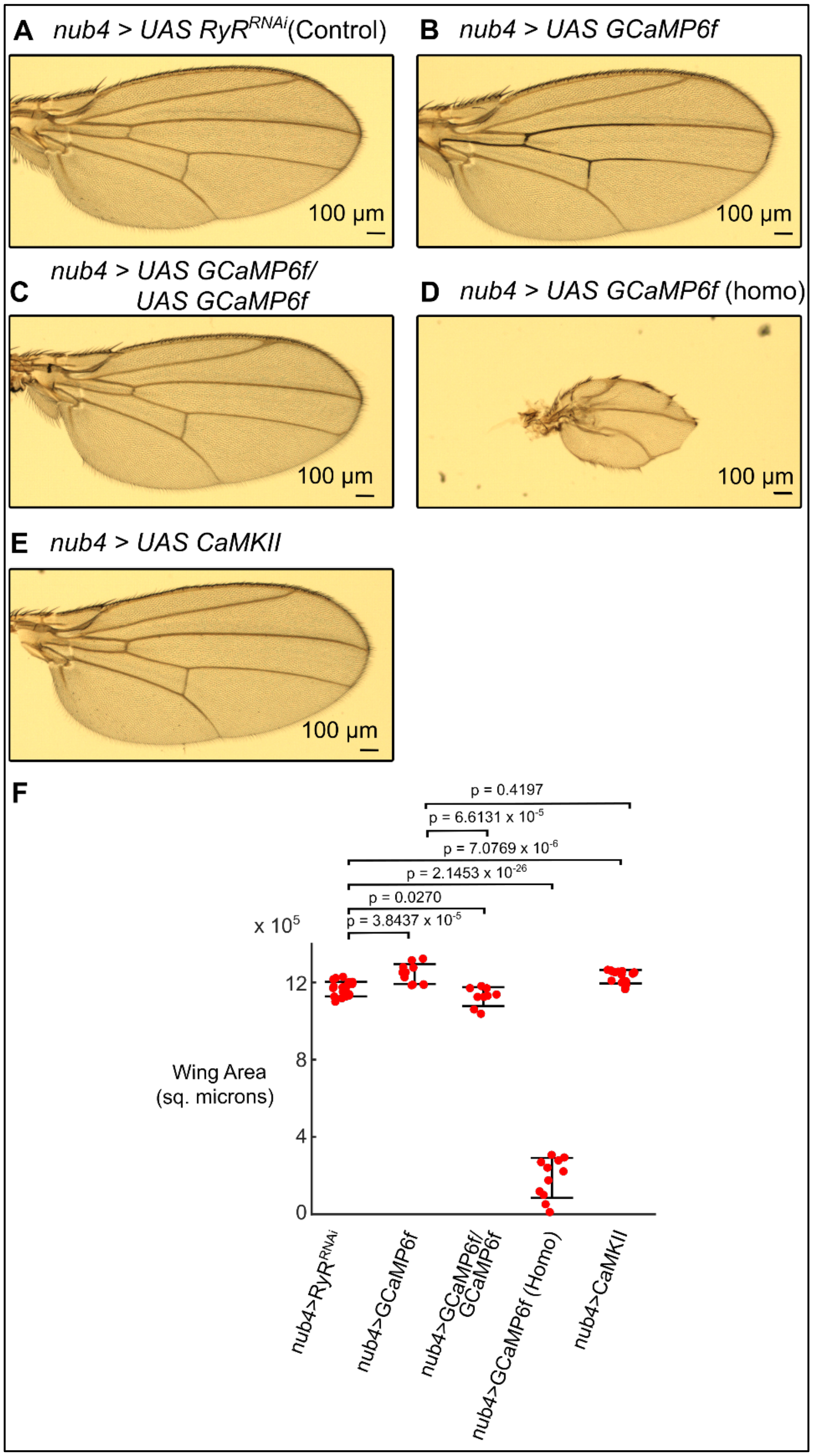


**Figure S8: Increasing gene dosage of GCaMP6f $Ca^{2+}$ sensor dramatically reduces wing size. (A-E)** Wings from adult males of indicated genotypes **(A)** nubG4, UAS $RyR^{RNAi}$, **(B)** nubG4, UAS GCaMP6f, **(C)** nubG4, UAS GCaMP6f/UAS GCaMP6f, **(D)** nubG4, UAS GCaMP6f (Homozygous) **(E)** nubG4, UAS CaMKII **(F)** Quantification of adult wing sizes. As the gene dose of GCaMPf is increased in the wing disc, the overall wing area decreases in size (B,C,D). overexpressing possible $Ca^{2+}$ downstream target CaMKII increases the wing size consistent with B in which one copy of GCaMP6f was expressed

**Table S1: Baseline parameters used in the model.**

| Parameter | Description | Value |
|---|---|---|
| $k_{5,P}$ | $IP_3$ degradation rate constant | 0.66 $s^{-1}$ |
| $K_{PLC}$ | Half activation of PLC | 0.2 μM |
| $V_{PLC}$ | Maximum production rate of PLC | 1.5 μM $s^{-1}$ |
| $\beta$ | Ratio of effective volumes ER/cytosol | 0.185 |
| $V_{SERCA}$ | Maximum SERCA pump rate | 0.9 μM $s^{-1}$ |
| $K_{SERCA}$ | Half activation constant | 0.1 μM |
| $k_1$ | Maximum rate of $Ca^{2+}$ release | 1.11 $s^{-1}$ |
| $k_2$ | $Ca^{2+}$ leak | 0.0203 $s^{-1}$ |
| $K_a$ | $Ca^{2+}$ binding to activating site | 0.08 μM |
| $K_i$ | $Ca^{2+}$ binding to inactivating site | 0.4 μM |
| $K_p$ | $IP_3$ binding | 0.13 μM |
| $\tau_{max}$ | Maximum time constant of $IP_3R$ inactivation | 800 $s^{-1}$ |
| $k_\tau$ | $Ca^{2+}$ dependent rate of $IP_3R$ inactivation | 1.5 μM |
| $F_p$ | GJ permeability for $IP_3$ | 0.005 $\mu M^2 s^{-1}$ |
| $F_c$ | GJ permeability for $Ca^{2+}$ | 0.0005 $\mu M^2 s^{-1}$ |

Most baseline parameters were adopted from Politi and colleagues (Politi et al., 2006).

**Table S2: Extended data movies**

| SI Movie # | Description |
|---|---|
| 1 | *nub-Gal4>UAS-GCaMP6f*, *UAS-mcherry*, ex vivo, spike |
| 2 | *nub-Gal4>UAS-GCaMP6f*, *UAS-mcherry*, ex vivo, ICT |
| 3 | *nub-Gal4>UAS-GCaMP6f*, ex vivo, ICW |
| 4 | *nub-Gal4>UAS-GCaMP6f*, ex vivo, fluttering |
| 5 | *nub-Gal4>UAS-GCaMP6f*, in vivo, spikes |
| 6 | *nub-Gal4>UAS-GCaMP6f*, in vivo, ICT |
| 7 | *nub-Gal4>UAS-GCaMP6f*, in vivo, ICW |
| 8 | *nub-Gal4>UAS-GCaMP6f*, in vivo, fluttering |
| 9 | Spike, Simulation output |
| 10 | ICT, Simulation output |
| 11 | ICW, Simulation output |
| 12 | fluttering, Simulation output |
| 13 | *nub-Gal4>UAS-GCaMP6f*, ex vivo in Grace's low 20E media, gap junctions not blocked (Control) |
| 14 | *nub-Gal4>UAS-GCaMP6f*, ex vivo in Grace's low 20E media with Carbenoxolone, gap junctions blocked |
| 15 | *nub-Gal4>UAS-GCaMP6f*, *UAS-RyR$^{RNAi}$*, ex vivo in Grace's low 20E media (Control) |
| 16 | *nub-Gal4>UAS-GCaMP6f*, *UAS-InsR$^{CA}$*, ex vivo in Grace's low 20E media |
| 17 | *nub-Gal4>UAS-GCaMP6f*, *UAS-InsR$^{DN}$*, ex vivo in Grace's low 20E media |
| 18 | *nub-Gal4>UAS-GCaMP6f*, *UAS-Gaq$^{OE}$*, ex vivo in Grace's low 20E media |

**Table S3:** Changes in wing area for known perturbations through GAL4/UAS system. Note that maximal deviations wing size for strong growth perturbations is in range of 20-50%.

| Pathway | Perturbation | Genotypes of perturbations | Genotypes of Control | % Changes in wing area | References |
|---|---|---|---|---|---|
| Insulin | InsR$^{CA}$ (Upregulation) | *nubGal4>UAS-GCaMP6f, UAS-InsRCA* | *nubGal4>UAS-GCaMP6f, UAS-mcherry* | 29% | This study |
| | InsR$^{DN}$ (Downregulation) | *nubGal4>UAS-GCaMP6f, UAS-InsRDN* | *nubGal4>UAS-GCaMP6f, UAS-mcherry* | -49% | |
| Ca$^{2+}$ | Gαq$^{OE}$ | *MS1096Gal4 > UAS-GαqOE* | *MS1096Gal4 > UAS-RyRRNAi* | -20% | (Brodskiy et al., 2019) and this study |
| | Gαq$^{RNAi}$ | *MS1096Gal4 > UAS-GαqRNAi* | *MS1096Gal4 > UAS-RyR*$^{RNAi}$ | -17% | |
| | itp-83A$^{RNAi}$ | *MS1096Gal4 > UAS-itpr-83ARNAi* | *MS1096Gal4 > UAS-RyRRNAi* | -11% | |
| | inx2$^{RNAi}$ | *MS1096Gal4 > UAS-inx2RNAi* | *MS1096Gal4 > UAS-RyRRNAi* | -29% | |
| | sl$^{RNAi}$ | *MS1096Gal4 > UAS-slRNAi* | *MS1096Gal4 > UAS-RyRRNAi* | -13% | |
| | plc21c$^{RNAi}$ | *MS1096Gal4 > UAS-plc21CRNAi* | *MS1096Gal4 > UAS-RyRRNAi* | -3% | |
| Hippo | exRNAi | *nubGal4>UAS-exRNAi* | *nubGal4* | 60% | (Su et al., 2017) |

| | | | | | |
|---|---|---|---|---|---|
| | Kib | *nubGal4>UAS-kib* | *nubGal4* | -20% | |
| Mechanical | $rok^{RNAi}$ | *Nub-Gal4>UAS-dcr2, UAS-rok*$^{RNAi}$ | *nubGal4>UAS-dcr2* | -20% | (Rauskolb et al., 2014) |
| | rok.CAT | *nubGal4>UAS-dcr2, UAS-rokCAT* | *nubGal4>UAS-dcr2* | 12% | |
